# A novel open-source cultivation system helps establish the first full cycle chemosynthetic symbiosis model system involving the giant ciliate *Zoothamnium niveum*

**DOI:** 10.1101/2024.11.03.621766

**Authors:** PE Contarini, E Emboule, P Jean-Louis, T Woyke, SV Date, O Gros, J-M Volland

**Author notes:** corresponding author: <u>Jean-Marie Volland</u>.

## Abstract

Symbiotic interactions drive species evolution, with nutritional symbioses playing vital roles across ecosystems. Chemosynthetic symbioses are globally distributed and ecologically significant, yet the lack of model systems has hindered research progress. The giant ciliate *Zoothamnium niveum* and its sulfur-oxidizing symbionts represent the only known chemosynthetic symbiosis with a short life span that has been transiently cultivated in the laboratory. While it is experimentally tractable and presents a promising model system, it currently lacks an open-source, simple, and standardized cultivation setup. Following the FABricated Ecosystems (EcoFABs) model, we leveraged 3D printing and polydimethylsiloxane (PDMS) casting to develop simple flow-through cultivation chambers that can be produced and adopted by any laboratory. The streamlined manufacturing process reduces production time by 86% and cuts cost by tenfold compared to the previous system. Benchmarking using previously established optimal growth conditions, the new open-source cultivation system proves stable, efficient, more autonomous, and promotes a more prolific growth of the symbiosis. For the first time, starting from single cells, we successfully cultivated the symbiosis in flow-through chambers for 20 days, spanning multiple generations of colonies that remained symbiotic. They were transferred from chamber to chamber enabling long-term cultivation and eliminating the need for continuous field sampling. The chambers, optimized for live imaging, allowed detailed observation of the synchronized growth between the host and symbiont. Highlighting the benefit of this new system, we here describe a new step in the first hours of development where the host pauses growth, expels a coat, before resuming growth, hinting at a putative symbiont selection mechanism early in the colony life cycle. With this simple, open-source, cultivation setup, *Z. niveum* holds promises for comparative studies, standardization of research and wide adoption by the symbiosis research community.

## Introduction

Symbiotic interactions, defined as associations between two or more organisms regardless of benefit, are commonly observed throughout the tree of life. Such interactions are ubiquitous, diverse and remain powerful drivers of evolution. Despite their ubiquity, fundamental mechanisms that lead to establishment of symbiotic interactions, their maintenance and even their breakdown are not very well understood. Species establish symbiotic interactions primarily to access more nutrients or diversify their diet (Douglas, 1998; Cavanaugh et al., 2006; Mizrahi, 2013; Fronk and Sachs, 2022). Based on observed associations, nutritional symbioses can be classified into three types: a) Heterotrophic, where symbionts feed on organic matter, *e.g. Epulopiscium* spp. (Sannino et al., 2023), represented by model systems such as the cow rumen (Wolin, 1979; Sharp et al., 1998), aphids (Brisson and Stern, 2006) and termites (Brune, 2014); b) Phototrophic, where symbionts obtain carbon produced through photosynthesis (Venn et al., 2008), represented by established model systems such as the green *Hydra* (Kovacevic, 2012), *Aiptasia* sp. (Rädecker et al., 2018) or the emerging system *Cassiopeia* sp. (Rädecker et al., 2017); and c) Chemosynthetic, where bacterial symbionts fix inorganic carbon/nitrogen using the energy gained from the oxidation of reduced inorganic compounds (Dubilier et al., 2008; König et al., 2017; Petersen et al., 2016). While multiple model systems for phototrophic and heterotrophic symbioses have been established, model systems for chemosynthetic symbioses, where symbionts maintain stable associations over several cycles of growth, are not available to the scientific community.

Here we describe successful cultivation of *Zoothamnium niveum*, a giant colonial ciliate characterized by a symbiotic relationship with the chemosynthetic sulfur-oxidizing bacteria *Candidatus* Thiobius zoothamnicola, using a novel, 3D-printed, flow-through culture system. Thiobius ectosymbionts, which form a monolayer covering entire *Z. niveum* colonies except for the basal part of the stalk (Rinke et al., 2006; Bright et al., 2014), serve as a nutritional source for the ciliate by providing essential nutrients through chemosynthesis (Volland et al. 2018). The giant ciliate stands out as a promising model system due to several key factors: it is the sole chemosynthetic model organism cultivable in a complete cycle in laboratory conditions with a short reproductive cycle; its considerable size makes it easy to manage in laboratory settings; and its simplicity with only two partners, one host, and one symbiont, facilitates experimental investigations (Bright et al., 2014; Volland et al., 2018). The symbionts are vertically transmitted as they adhere to the surface of macrozooids which eventually leave the parent colony as swarmers to establish new colonies (Bauer-Nebelsick et al., 1996a, 1996b). *Zoothamnium niveum* demonstrates asexual reproduction *in vitro*. Our cultivation likely represents the first model system for chemosynthetic symbiosis (Herron et al., 2013; Corliss, 2002) that is stable and which can be manipulated under laboratory conditions.

*Zoothamnium niveum* colonies inhabit the interface between anoxic, sulfidic environments and oxic, non-sulfidic environments, where they thrive due to their unique physiological adaptations (Espada-Hinojosa et al., 2022). In order to successfully grow *Z. niveum* in laboratory conditions, it must be provided with hydrogen sulfides (symbiont’s electron donor) and oxygen (host’ and symbiont’s electron acceptor) to meet the oxygen demand of both the symbiont and the host. Because oxygen rapidly oxidizes sulfides abiotically a stable liquid media cannot be prepared in a closed system. To address this, a flow-through culture system that continuously mixes oxic seawater and anoxic hydrogen sulfide in custom-made chambers was designed in 2007 and allowed a first cultivation of the symbiosis using entire colonies sampled from the environment as an inoculum (Rinke et al., 2007). Later, these chambers were redesigned and used to grow colonies from single-cells (symbiotic and aposymbiotic) and highlighted symbiont-induced polyphenism in the ciliate host (Bright et al., 2019). Thus, *in vitro* cultivation of *Zoothamnium niveum* has been achieved, but only for limited duration (13 days for Rinke et al. 2007; 7 days for Bright et al., 2019) and only for a single generation of adult colonies. Additionally, there was no transfer of colonies between chambers, maintaining dependence on field collections for new cultures. These studies marked progress in *Z. niveum* cultivation but fell short of establishing sustained, independent laboratory cultures with multiple generations. There was also a mandatory need for daily maintenance and the cultivation system used by Bright et al. (2014) is neither commercially available nor open-source and therefore not more broadly available to the scientific community.

Establishing symbiotic models systems has traditionally been difficult as it involves cultivating two or more species in an in-vitro setting under strict conditions that mimic complex environments. Our newly developed experimental setup accomplishes this goal (Duffy et al., 2021). The system passes all benchmarks pertaining to tractability, stability and reproducibility of model systems, successfully maintaining various chemical conditions needed for survival and proliferation of *Z. niveum* in the presence of its symbiont. Our 3D-printing approach largely overcomes the limitations of previous setups, such as high costs, challenges in reproducibility, reliance on external manufacturing, and intensive maintenance requirements. Colonies of Z. niveum cultivated in Guadeloupe were compared with colonies collected and analyzed from California, allowing us to confirm the presence of the same host-symbiont association across different geographic locations, further supporting the robustness of this system as a model for chemosynthetic symbiosis. Our goal is to provide the scientific community with an open-source, easy to access and easy-to-reproduce, straightforward cultivation system that is capable of maintaining stable symbiotic associations between species over multiple generations. Our approach also makes it feasible to attempt cultivation of other chemosynthetic symbioses under laboratory conditions.

## Materials and methods

### Sampling

Colonies of the ciliate *Zoothamnium niveum* were primarily collected in the marine part of the mangrove from the “Manche-à-eau” lagoon (N-16.27’62”49, W-61°55’53”84), Guadeloupe (Lesser Antilles) at approximately 1.30 m depth, on organic substrates (mainly leaves and small branches of the mangrove tree *Rhizophora mangle*). Colonies from a tide pool at White Point Beach (San Pedro, California) were collected and observed under a dissection microscope, identified as putative *Z. niveum* colonies based on their morphology and bright white appearance, and flash frozen in liquid nitrogen until further processing.

### DNA preparation, sequencing and genome analysis

Six frozen *Zoothamnium niveum* colonies from White Point Beach, California were thawed and used for genomic DNA extraction using the ZymoBIOMICS DNA Microprep Kit (Zymo Research, Irvine, CA). The manufacturer’s protocol was followed, with the following modifications: the colonies were added to a ZR BashingBead Lysis Tube (0.1 & 0.5 mm beads) with 750 µl ZymoBIOMICS Lysis Solution. The sample was processed with a Biospec Mini-Beadbeater (Biospec, Bartlesville, OK) for 1 minute at the maximum “homogenize” setting before transfer into a Zymo-Spin III-F Filter in a collection tube. The manufacturer’s protocol was followed thereafter and the final DNA extract was eluted in 25 µl from a Zymo-Spin IC Column. DNA libraries were created from 1ng of DNA using Nextera XT DNA library creation kit (Illumina). We sequenced the DNA libraries at the DOE Joint Genome Institute on an Illumina NovaSeq S4 platform. We then imported pair-end reads (2×150 bp) into the KBase platform (www.kbase.us) (Arkin et al 2018) where we used MetaSPAdes (v3.15.3) to assemble reads into contigs of at least 2000 bp (using kmers of 33, 67, 99, 125 bp). We then binned contigs using MetaBAT2 (v1.7) resulting in 7 bins (Supplementary Table 1). One of the bins was taxonomically identified as a Chromatiales by the GTDB-Tk classify app (v1.7.0), which corresponded to the symbiont *Ca*. Thiobius zoothamnicola. This bin was extracted and treated as a genome assembly for further analyses (referred to as *Ca*. T. zoothamnicola Metagenome Assembled Genome (MAG) or symbiont MAG). The symbiont MAG was annotated with Prokka (v1.14.5) and we assessed genome quality with CheckM (v1.0.18) (Table S3). We built a phylogenetic tree of the symbiont MAG assembly and 24 related gammaproteobacterial genomes available in public databases using SpeciesTreeBuilder (v0.1.4) in Kbase. To build the tree we used a set of 49 core, universal genes defined by COG (Clusters of Orthologous Groups) gene families (COG0012, AlaS, PheS, ArgS, KsgA, PurE, PurL, RpsL, RpsG, RpsJ, RpsB, PheT, RplK, RplA, AroC, RpoC, RplC, RplD, RplW, RplB, RplV, RpsC, RplN, RplE, RpsH, RplF, RpsE, RpsM, RpsK, RplM, RpsI, Ndk, Pgk, COG0127, TruB, PurM, PurD, RnhB, SerS, RpsS, RpsQ, CysS, RplJ, RplR, Tgt, PyrG, GuaA, InfB, QRI7). We computed pairwise Average Nucleotide Identities (ANIs) with FastANI (v0.1.3). The filtered raw reads data of the metagenome can be accessed from IMG with the ID number 3300063003. The complete metagenome assembly, the seven bins, including the symbiont MAG, and their respective annotations can be accessed from the Kbase narrative named “*Zoothamnium niveum* metagenome from California (public)” at : https://narrative.kbase.us/narrative/190725.

### Preparation of the cultivation chamber

A polydimethylsiloxane (PDMS; SYLGARD 184®) 10:1 mixture base to the curing agent is degassed in a vacuum chamber for 30 minutes and poured into the assembled printed mold of the chamber. The mold is heated at 50°C overnight. After solidification, the PDMS is carefully removed from the mold and trimmed if needed. The PDMS chambers were then soaked in 95% ethanol overnight, the next day they were rinsed with distilled water and allowed to air dry.

The cultivation chamber consists of a polydimethylsiloxane (PDMS) body (internal dimension 15mm X 35mm X 10mm), a standard glass slide (51 mm by 75 mm), and a 3D-printed frame that ensures the glass slide is sealed onto the PDMS body. The frames are held together by six bolts and nuts (Fig 1). When assembled, these components form the cultivation chamber. To connect the cultivation chamber to the static mixer (FRIGIIRE® Mixing Nozzles 3.6 in, 1:1 & 2:1 ratios) and the flow-through system, 15 Gauge blunt needles are sharpened with fine grain sand paper and carefully inserted straight into the PDMS on both ends of the device. The technical drawings for the central part and the side of the mold are provided in supplementary figures S1 and S2 respectively. The technical drawing for the frame is provided in supplementary figure S3. The 3D model for all printed components are available for download at : https://github.com/jvolland/Zoothamnium-Cultivation-System.

**Figure 1.**
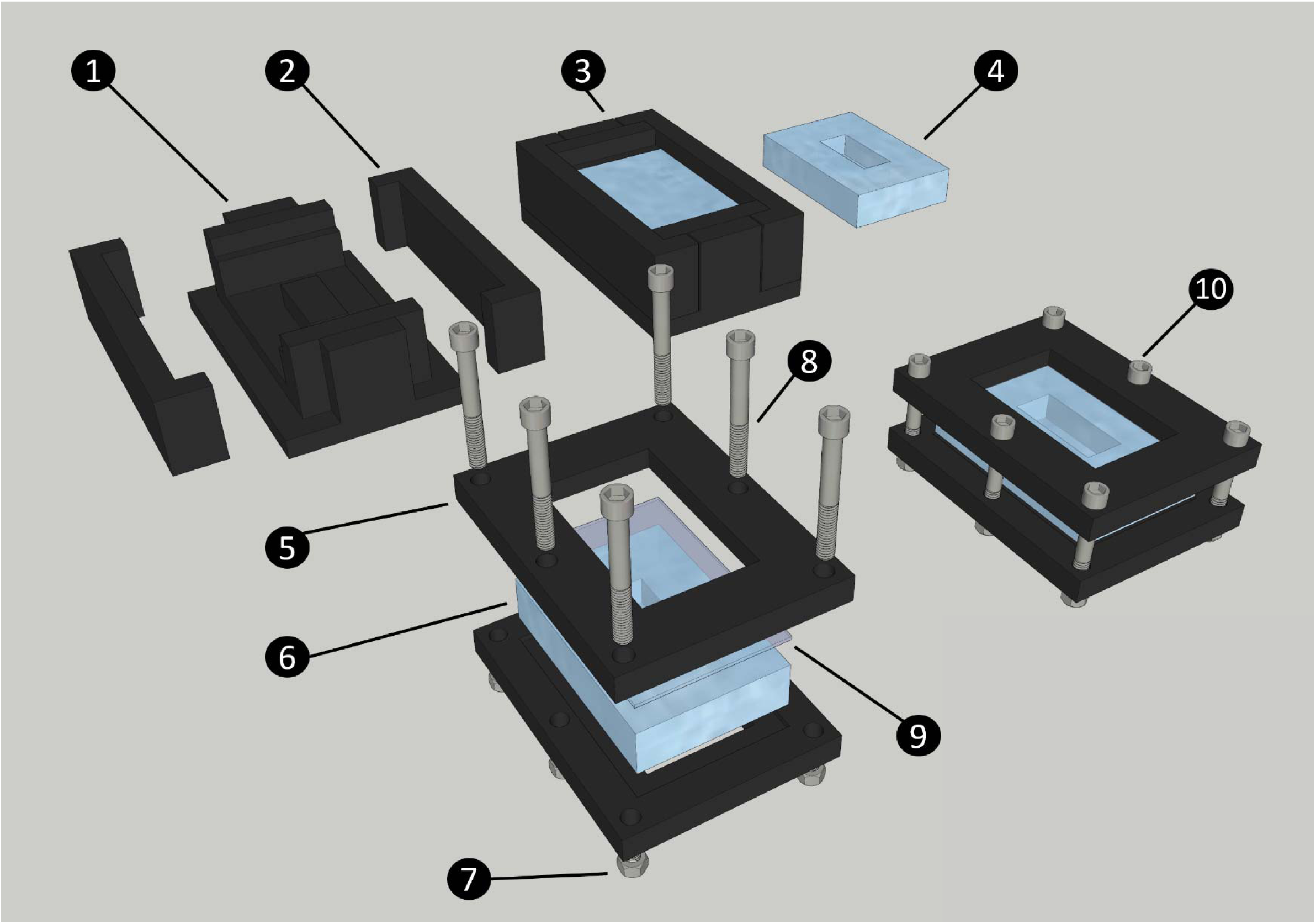
Diagram of the steps to produce and assemble the cultivation chamber. The chamber is made of a central PDMS body, closed by a standard glass slide (75 x 51 mm). Both these parts are maintained together by two 3D printed frames secured with nuts and bolts. The 3D printed chamber mold is made of a central part (1) and two sides (2); The liquid PDMS is poured into the closed mold and secured with high-temperature polypropylene plastic tape (not shown on diagram) to form the chamber (3); PDMS Chamber removed from mold after curing (4); Printed chamber frame (5); PDMS chamber (6); Nut (7); Bolt (8); Glass slide (9); Assembled cultivation chamber (10).

### Hydrogen sulfide and filtered seawater preparation

The hydrogen sulfide solutions used for the cultivation are made from Na_2_S.9H_2_O. An anoxic stock solution was prepared at a concentration of 50 mM in a crimp bottle closed by a thick rubber septum and an aluminum cap. This solution of distilled water was bubbled with argon gas for 30 minutes, before adding Na_2_S.9H_2_O crystals for a final concentration of 50 mM. This anoxic concentrated stock can be stored at 4℃ for several weeks.

The stock sulfide solution was then diluted to prepare a final sulfide solution with NaCl at 20 g.L^-1^. Addition of NaCl improves the mixing of the sulfide solution with seawater by matching its viscosity. To prepare the final sulfide solution, saline MilliQ water was bubbled with argon gas for 30 minutes before adding the stock sulfide solution to get a concentration of 0.5 mM inside the final sulfide solution. This final sulfide solution was prepared just before use and replaced twice a week. The seawater used for cultures was filtered at 1.2 μm to reduce the microbial load and exclude the meiofauna.

The filtered sea water and the final sulfide solution continuously mix in the static mixer to achieve the required operational concentration in the culture chambers, between 3 and 33 µmol/L (Rinke et al., 2007).

### Culturing in chambers

Colonies retrieved from the environment were cut at the base of the stalk and transferred to an embryo dish filled with normoxic 0.2 μm filtered seawater to induce the release of swarmer (as described in Espada-Hinojosa et al., 2022). The swarmers were collected using a Pasteur pipette and transferred into the culture chamber. To induce the fixation of the swarmers, a high concentration of hydrogen sulfide was added directly into the chamber in order to obtain a theoretical final concentration of 0.3 mM. The closed chamber was left untouched for approximately 5 hours giving the swarmers time to settle onto the walls of the chamber. The chamber was then plugged in a continuous flow of, with a regular supply of hydrogen sulfide, ensuring the necessary conditions for the swarmers to settle and thrive.

### Continuous flow system

The system consists of two Ismatec® peristaltic pumps: one delivers 1.2 µm-filtered normoxic seawater, while the other supplies an anoxic hydrogen sulfide solution prepared in distilled water with 20 g.L of NaCl to match the viscosity of seawater. The seawater and the sulfide solution are homogeneously mixed by a helical static mixer before entering the culture chamber. At the outlet of the chamber, a tubing routes the flow into a waste container (Fig 2 and Supplementary figure S4). By mixing the seawater:sulfide by using a 10:1 the output flow rate is established at 60 mL/h.

**Figure 2.**
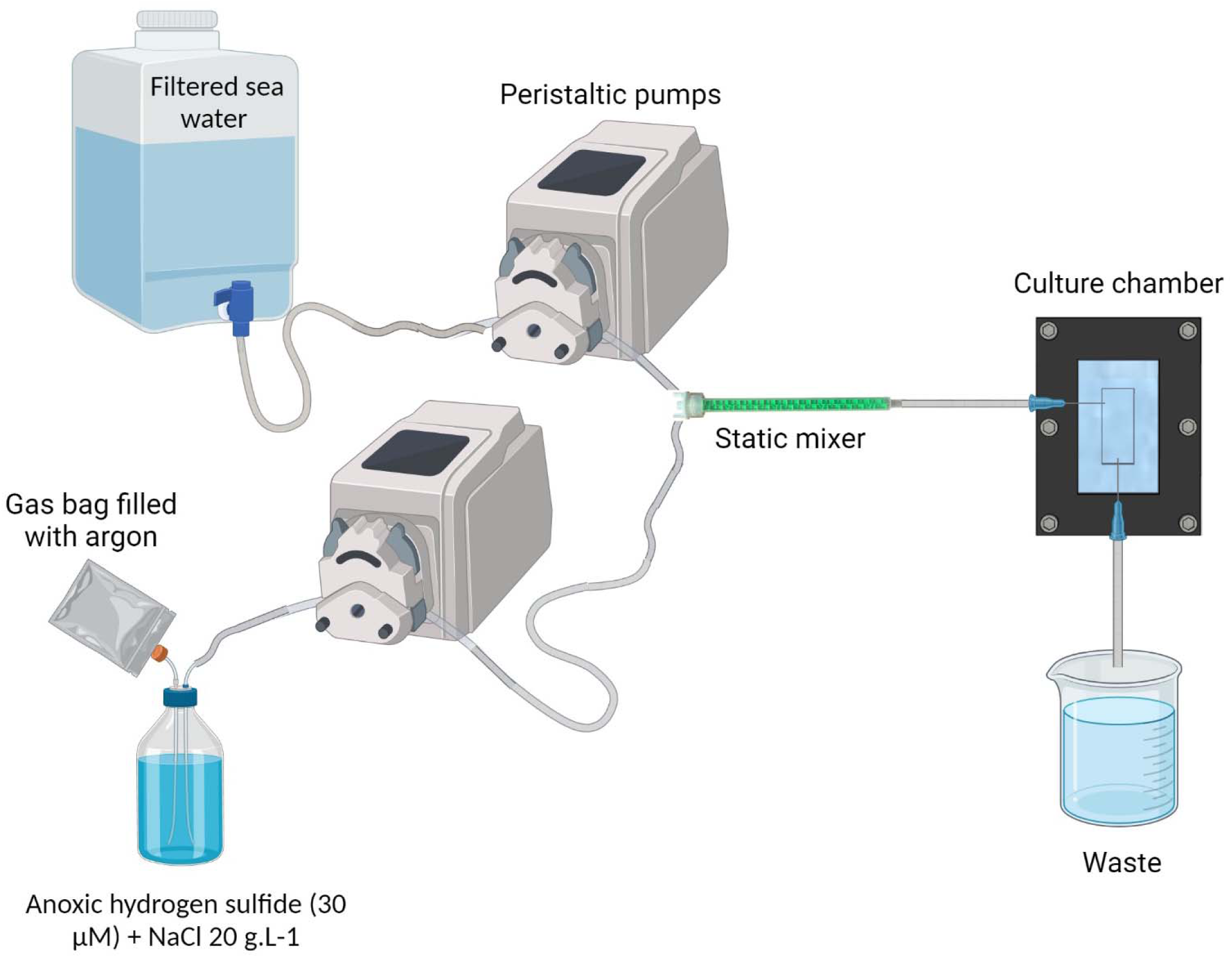
Complete cultivation system. NaCl is added into the anoxic hydrogen sulfide solution, at 20 g/L concentration. A gas bag filled with argon provides a reserve of anoxic gas to replace the sulfide solution in the bottle and prevents the remaining sulfide from being oxidized. The filtered sea water and the anoxic hydrogen sulfide water are pumped through peristaltic pumps and meet inside the static mixer to properly mix the fluids. The resulting oxic-sulfidic sea water enters the culture chamber, providing the ciliate’s ectosymbionts with both electron donor and acceptor. Created in BioRender. Contarini, P. (2023) BioRender.com/h94n742.

### Monitoring

Throughout each *Z. niveum* cultivation cycle, daily monitoring of physico-chemical parameters was conducted to ensure optimal growth conditions. These parameters included temperature, salinity, pH, flow rate, and sulfide concentration in the effluent water. Temperature measurements were obtained using a thermometer, salinity levels were assessed using an Atage S-10 refractometer, and pH values were determined using a Mettler Toledo© pH probe. To measure the flow rate, water was sampled from the system outlet over a 30-minute period and the flow rate was then expressed in milliliters per hour. Sulfide concentrations were quantified using Cline’s method as detailed in Reese et al. (2011). Microbial abundance in cultures #3 and #4 was assessed on days 7, 15, and 20 by fixing 1 mL samples in 4% paraformaldehyde. Samples were filtered through 0.2 µm black polycarbonate Nuclepore membranes, stained with SybrGreen, and analyzed under a Leica TCS SPE confocal microscope.

Quantitative analysis of bacterial counts was performed by evaluating ten random 120 µm² fields per sample, and statistical differences between sampling days were analyzed using ANOVA.

### Growth monitoring

Growth monitoring was carried out daily in each culture and the timelapse of the beginning of the development of a colony of *Z. niveum* was done by taking a photo every 15 seconds for 8 hours. This monitoring was carried out using a dissecting microscope (Wild M3C Heerbrugg Switzerland) equipped with a Canon EOS 700D camera. Each colony was photographed every day in order to obtain the number of colonies within the chamber, and the number of branches per colony. More occasionally, timelapses of growing newly fixed colonies or of colonies’ contractions were made.

### Scanning electron microscopy

The control of the presence and the morphology of the symbionts were observed under a scanning electron microscope on colonies of generations G0, G1, and G2. These colonies were fixed in a solution of 2.5% glutaraldehyde in 0.2um filtered seawater for at least 2h at 4°C and rinsed twice with filtered seawater. They were then dehydrated in a graded acetone series, critical point dried with CO_2_ as transitional fluid (EM CPD300, Leica), and sputter-coated with gold (Sputter coater SC500, Bio Rad) before observation using an FEI Quanta 250 microscope at 20 kV.

## Results

### Zoothamnium niveum sampling sites

In addition to the 8 known sampling sites, 7 new sampling sites were identified through a search of the scientific literature, of taxonomic inventories, and through exploration of new locations along the Pacific coast of the USA (Fig 3; Supplementary table 1). Large white ciliate colonies from the Pacific coast of the USA (White Point Beach California), morphologically identical to *Zoothamnium niveum*, were analyzed by metagenomics. Subsequent sequencing of the colonies confirmed their identity.

**Figure 3.**
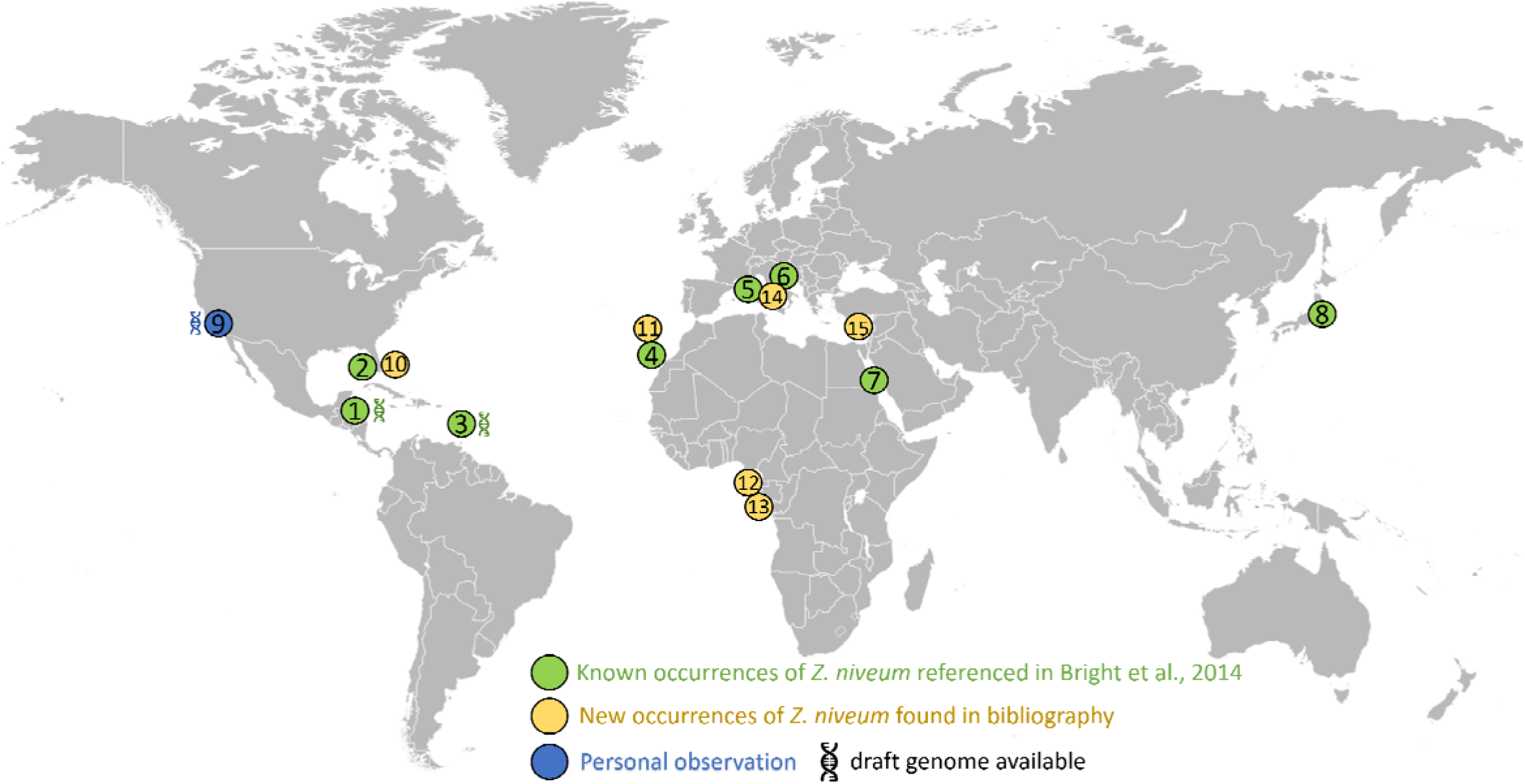
Distribution of the documented presence of *Zoothamnium niveum*. The listed sites span diverse coastal locations worldwide. These include: Twin Cays Island in Belize has an available genome for *Thiobius zoothamnicola* Espada-Hinojosa et al., 2024 (1); Gulf of Mexico, near the Florida Keys, USA (2); Guadeloupe in the French West Indies available genome for *Thiobius zoothamnicola* Espada-Hinojosa et al., 2024 (3); Lanzarote located in the Canary Islands, Spain (4); Corsica, France (5); Adriatic Sea (6); Red sea (7); Tokyo bay (8); White Point Beach in California, USA available genome for *Thiobius zoothamnicola,* this paper (9); Indian River Lagoon as well as Pete stone’s creek located in Florida, USA (10); Madeira Island (11); São Tomé Island (12); Gabon (13); Giglio Island, Italy (14); and Cyprus Island (15). Supplementary table 1 provides more details on the *Z. niveum* locations, including literature citations.

### Genomic analysis

Metagenomics and phylogenetic analysis confirmed the taxonomic identification of the White Point Beach colonies as *Z. niveum*. A metagenome obtained from six colonies was sequenced and assembled. Using MetaBAT2 for contig binning and GTDBtk for taxonomic assignment we recovered a MAG for *Ca*. Thiobius zoothamnicola. Phylogenomic analysis positioned this MAG alongside the Guadeloupe *Z. niveum* symbiont (Espada-Hinojosa et al., 2024) (Fig 4). The *Ca*. T. zoothamnicola MAG from California is estimated to be 85.5% complete and comprised 1664 predicted genes. While slightly less complete, it has otherwise similar genome statistics compared to the MAG obtained from colonies sampled in the Caribbean (*i.e*. Belize and Guadeloupe)(Supplementary table 2). The California and Guadeloupe MAGs have an Average Nucleotide Identity (ANI) of 96.7%, exceeding the species threshold of 95% for ANI (Konstantinidis and Tiedje, 2005; Rodriguez-R L.M. et al., 2021). The 16S rRNA gene sequence of *Ca* Thiobius zoothamnicola from the metagenome of White Point Beach colonies is 99.7% identical to those from the Mediterranean (France: Bay of Calvi, Corsica, Rinke et al. 2007) (Supplementary data 1). The phylogeny, ANI and the 16S rRNA gene sequence confirmed that the symbionts from California and Guadeloupe belong to the same species. While the host genome was not assembled, we recovered a near complete 18S rRNA gene sequence from the colony’s metagenome from White Point Beach (Supplementary data 2). A nucleotide BLAST revealed *Zoothamnium niveum* as the top hit, with a 99.7% identity to the 18S rRNA gene of *Z. niveum* from Florida (Clamp and Williams, 2006).

**Figure 4.**
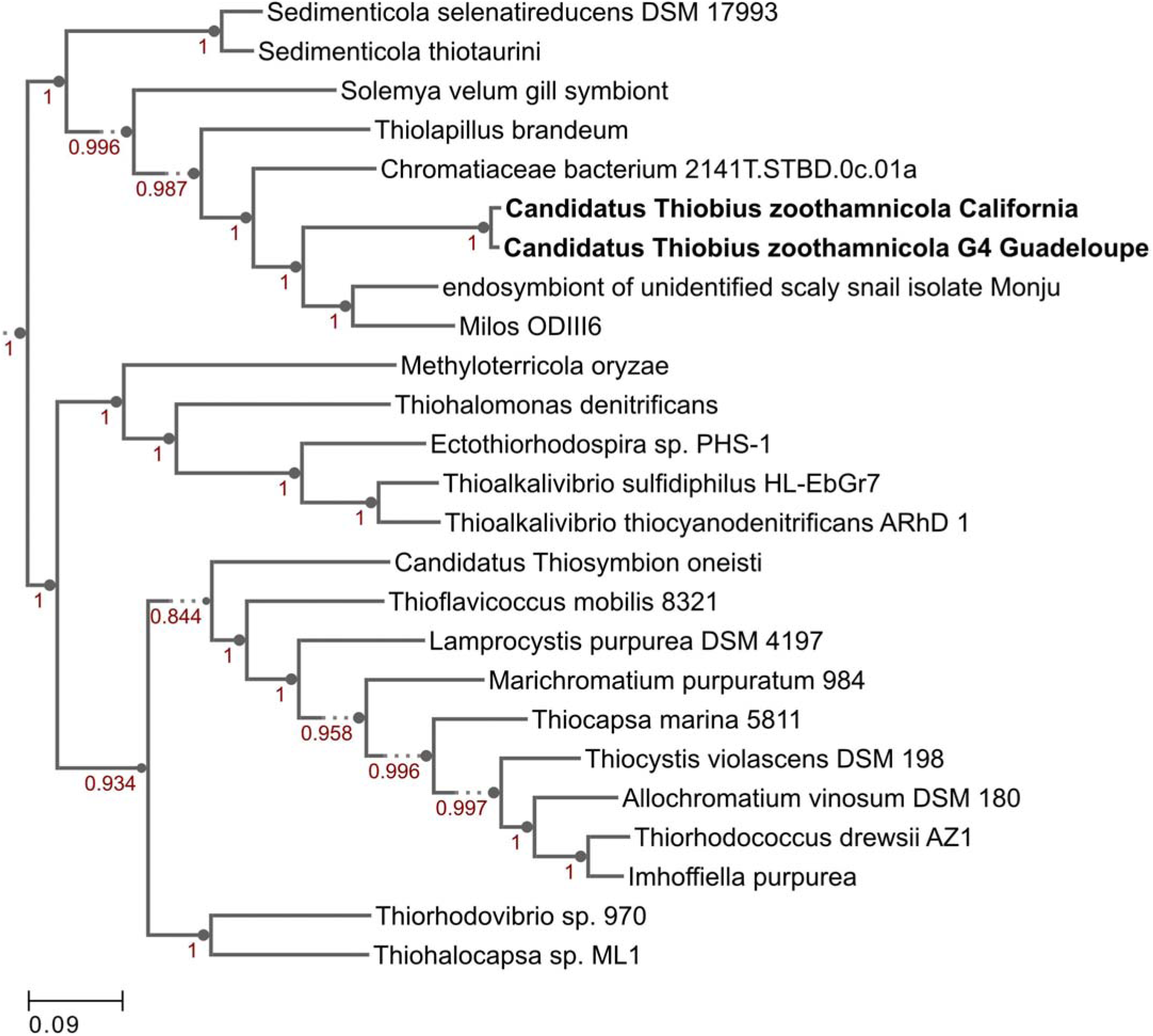
Genome phylogenetic tree of *Ca.* Thiobius zoothamnicola from California and Guadeloupe as well as 23 closely related Gammaproteobacteria. Maximum likelihood tree built using a set of 49 core, universal genes defined by COG gene families using SpeciesTreeBuilder (v0.1.4). Based on the symbiont genomes, it showed that the two *Ca*. T. zoothamnicola draft genomes from Guadeloupe and California are closely related.

### Cultivation system apparatus design

We simplified *Z. niveum* cultivation by designing an affordable and open-source cultivation system. It is composed of few parts that can be produced in any laboratory without the need of external manufacturing companies (table 1). Additionally, we implemented a large anaerobic sulfide reservoir equipped with a argon-filled gas bag, prolonging the anoxia of the H₂S solution. Furthermore, adjustment of the sulfide solution viscosity and the integration of a static mixer optimizes fluid mixing, addressing fluid stratification issues.

**Table 1.** Comparative table of the current and new culture chambers. For this new system, there is no need to hire an external company to make a custom chamber.

| Features | Current system | New system |
| --- | --- | --- |
| open source | no | yes |
| Number of parts | 25 | 16 |
| Manufacturing companies involved (no.) | 3 | 0 |
| Price per unit (approx.) | \$240 | \$20 |
| Approximate time of production (days) | 14 | 2 |
| Customizable geometry | No | Yes |

### Growth Monitoring

In order to demonstrate the efficacy of the new setup, we conducted four independent cultures which were stopped after 20 to 28 days. The data for culture #3 are shown on figure 5, while data for the three additional cultures are presented in supplementary data (respectively supplementary Figure S5, S6 and S7 for Culture 1, 2 and 4). To monitor the colonies throughout the growing period, photographs were taken at specific intervals. These photographs highlight the large number of colonies attached and developing, as well as the growth of the individual colonies (Fig 5A; Supplementary Figure S8). The physicochemical parameters were maintained in a stable state, with a continuous flow rate of 61.5 ± 3.32 mL/h, a pH of 7.48 ± 0.11, a temperature held at 24.9 ± 0.46 °C, a salinity of 34.9 ± 0.32 g/L, and a hydrogen sulfide concentration at 5.32 ± 0.09 µM (Fig 5B). The settlement rate was 41.5% with a total of 82 swarmers injected and 34 successfully fixed. With a max longevity of 17 days and an average colony lifespan of 11.90 ± 3.11 days. The average branches on day 13 was 67.8 ± 4 branches with a daily gain of 4.4 (±1) branches per day.

**Figure 5.**
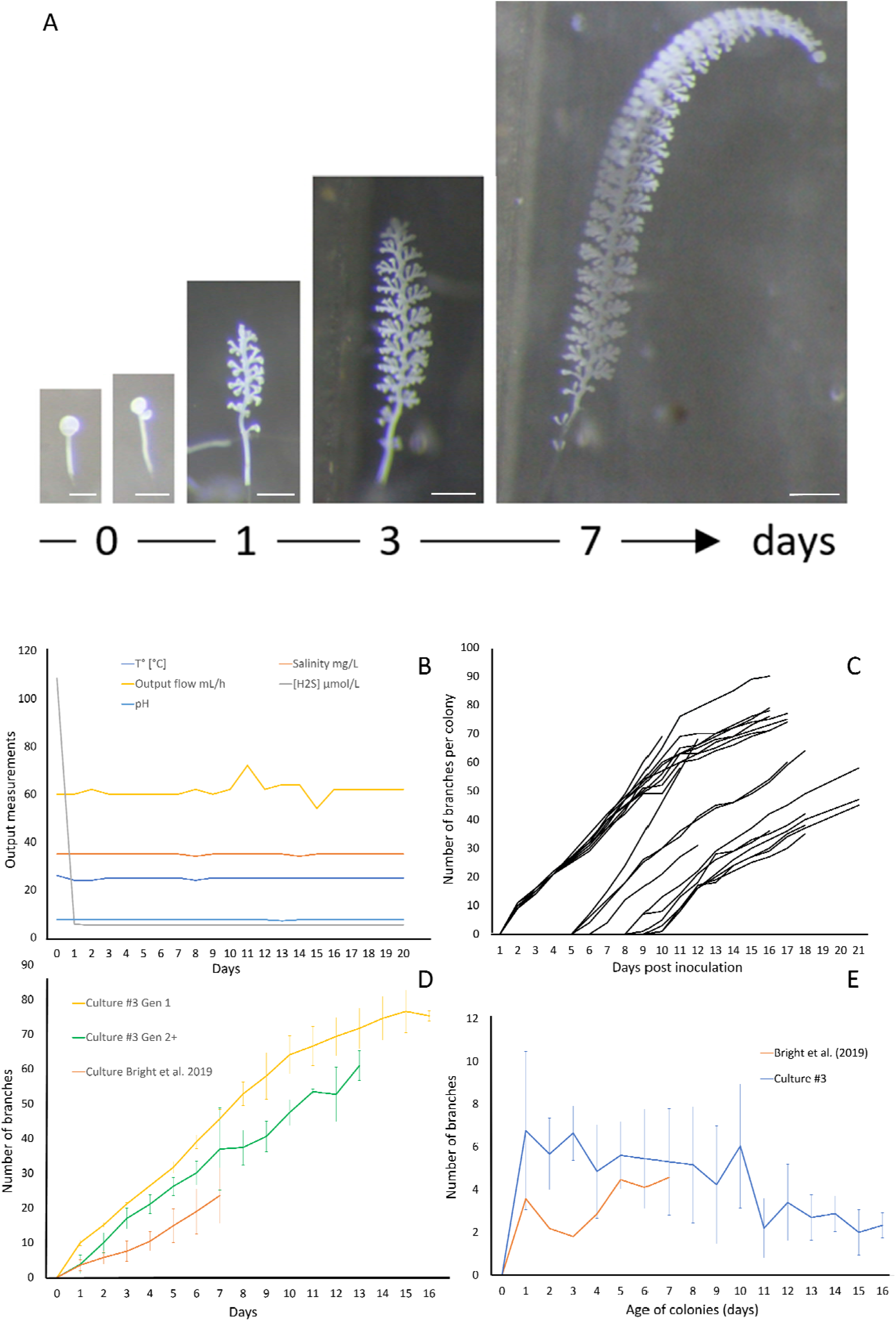
Growth of *Zoothamnium niveum* culture #3. (A) Example of the tracking of a single *Z. niveum* colony through a week (scale bar = 200 μm). Day 0, the swarmer has settled and begins the development of its stalk and its first divisions of the terminal zooid; Day 1, the stalk continues to develop, appearance of the first branches and the terminal zooid continues to divide; Day 3 and after, the stalk continues to grow and so does the number of branches. (B) Physico-chemical parameters measured in culture #3. To induce the attachment of the swarmers used as an inoculum, a higher sulfide concentration is established at day 0 and the chamber is left untouched without flow for several hours for the swarmers to settle. At the end of the settlement periode the flow is established and the normal sulfide concentration is maintained for the rest of the culture. ; (C) Growth of randomly selected individual colonies. The second generation of colonies start to grow after 5 days.; (D) Comparison of colony growth between the first generation (n=10), the following generations (n=11) and Bright et al. (2019; (E) Average branch gain per day measured on our culture (n=21) and Bright et al. (2019) (n=31).

The abundance of free-living microorganisms was monitored at day 7, 15 and 20 for cultures #3 and #4 (Supplementary Figure S9). The median microbial abundance (MA) was 4×10 in culture #3, and 2.6×10 in culture #4. The ANOVA statistical analysis showed no significant differences in microbial abundance between days 7, 15, and 20 in the two cultures examined. Stable environmental conditions in the culture chamber controlled the microbial abundance consistently across cultures, preventing the growth of free-living microbial organisms or biofilm that could either contaminate the culture chamber and overgrow the symbiosis.

The giant ciliate *Zoothamnium niveum* was successfully cultivated over multiple generations within each culture, which were intentionally terminated after approximately twenty days. After five days of cultivation (Fig 5C), a second generation of *Z. niveum* from the pioneer generation was obtained. It allowed us to compare the growth of colonies between generations (Fig 5D) as well as the average daily branch gain per colony within culture #3 (Fig 5E). The number of colonies increases continuously during the whole cultivation process in all cultures (Supplementary Figure S10).

We successfully transferred cultures from one chamber to another. Adult F2 colonies harboring macrozooids were collected after 8 days of in vitro culture and treated as colonies from mangrove to induce the release of swarmers outside of the chamber. These swarmers produced in vitro were transferred into a new culture chamber, settled successfully and subsequent generations were cultivated in the laboratory.

The presence of ectosymbionts on the surface of colonies was monitored by observing colonies under a scanning electron microscope. The wild colonies are called G0, whereas the first generation are referred to as G1 and the second generation as G2 (Fig 6). All generations, including G0, G1, and G2, were confirmed to consistently maintain their symbiotic relationship throughout the study.

**Figure 6.**
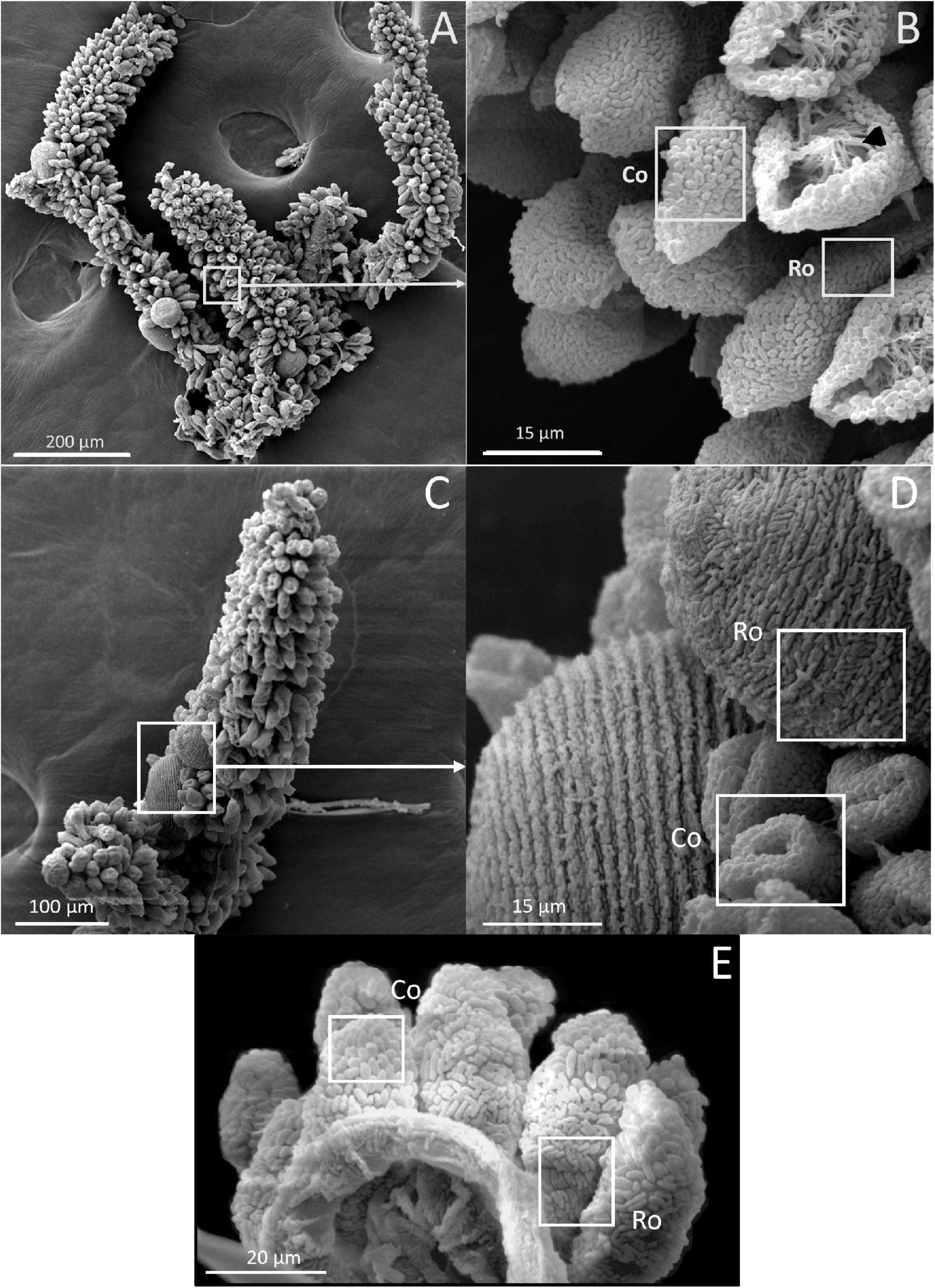
SEM images of *Z. niveum* colonies from the environment and from lab cultures. (A) Wild colony with macrozooids and microzooids retracted on the stalk; (B) zoom on that colony with two focus zones : first zone with microzooid covered with coccoid shaped symbionts (Co); base of a microzooid covered with rod shaped symbionts (Ro); black arrow showing the zooid’s oral ciliature. (C) observed F1 colony; (D) zoom on that colony; macrozooid covered with rod shaped symbionts (Ro); zooids covered with coccoid shaped symbionts (Co); (E) observed F2 colony with the focus zone; macrozooid covered with rod shaped symbionts (Ro); zooids covered with coccoid shaped symbionts (Co).

### Live imaging

To demonstrate the suitability of our setup for live imaging, we took time-lapse photographs of the first 10 hours of colony development in the chamber. We observed the growth of the stalk without symbionts, as the terminal zooid rotates on itself (Fig 7A). After a pause in growth, and the expulsion of a coat around the terminal zooid is noted (Fig 7B). Growth resumes after coat expulsion, followed by colonization of the rest of the colony by its symbionts, including the stalk (Fig 7C-H). A movie about the release of the coat is included as supplementary material (Supplementary movie 1). These coats were examined using both light and confocal microscopy after DNA labeling with SYBR green. Under light microscopy (Fig 7i), the coats appeared as distinct structures emerging from the zooid. They exhibited a spherical or ovoid structure with a ∼70µm radius. The surface of the coats was irregular and rough. Subsequent examination was performed using confocal microscopy (Fig 7J). DNA staining and CLSM reveal bacteria on coats with a pattern resembling terminal zooid symbionts. SYBR Green-marked bacteria share morphologies with symbionts, including rod and cocci forms. A timelapse shows the contraction movements of *Zoothamnium niveum*, a natural behavior of the host that facilitates the exchange of nutrients between it and its symbiont (Supplementary video 2).

**Figure 7.**
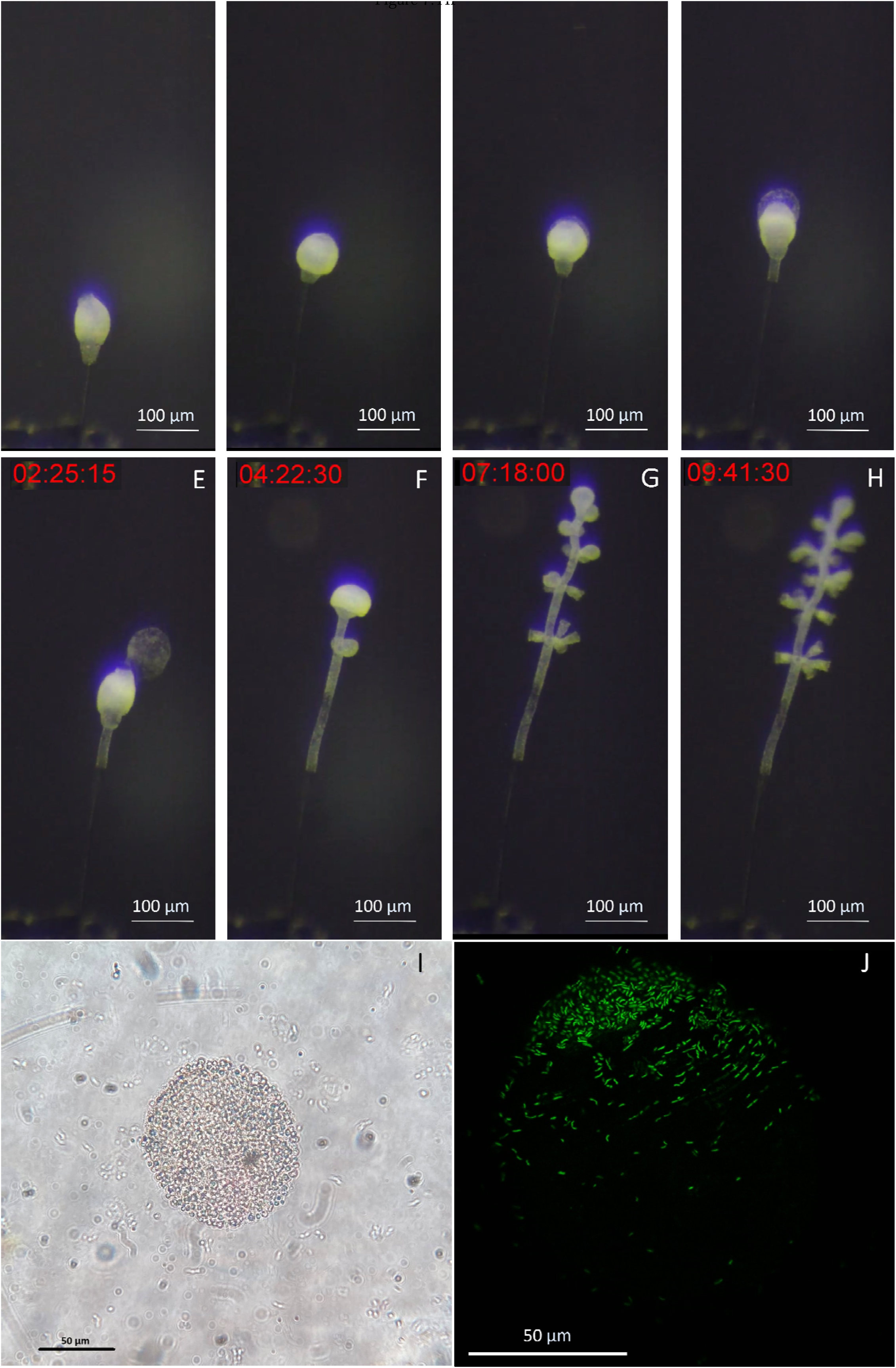
Images of the released terminal zooid’s coat. (A) The stalk grows without symbionts, as the terminal zooid rotates on itself. (B) There is a pause in growth, and the expulsion of a coat around the terminal zooid begins. (C and D) During the expulsion of the coat, growth resumes, this time with colonization of the rest of the colony by its symbionts, including the stalk; (E) The coat is expelled as the growth continues. (F, G and H) Microzooids are differentiated along the stem as the terminal zooid grows. (I) Light image of a coat taken from a zooid, observation of an ovoid structure with a ∼70µm-radius presenting roughness on its surface, suggesting bacteria; (J) Confocal image of a SYBR green staining of a coat taken from a zooid, observation of an ovoid structure with a ∼70µm-radius, presenting marked bacteria on part of its surface.

## Discussion

Chemosynthesis enables certain organisms to thrive in extreme environments with high concentrations of inorganic molecules like hydrogen sulfide, ferrous iron, hydrogen, or methane, found in deep sea environments (hydrothermal vents or cold seeps) or mangrove ecosystems (Dubillier et al., 2008). Recent studies suggest that chemosynthesis contributes more than previously thought to global carbon fixation (Ricci and Greening, 2024). This metabolism supports a diverse range of symbiotic relationships, showcasing adaptability in morphology among closely related host groups, particularly prevalent in ecosystems like mangroves, hydrothermal vents, anoxic basins, and cold seeps (Frenkiel et al., 1996; Petersen et al., 2011; Sogin et al., 2020). Chemosynthetic symbioses play an ecological role in sustaining complex ecosystems by forming the foundation of trophic networks that connect the lithosphere to the biosphere, supporting a wide range of organisms, including bacteria, archaea, small invertebrates belonging to the meiofauna, and sometimes even larger species such as tube worms and bivalves (Minic and Hervé, 2004; Roeselers and Newton, 2012). However, research on chemosynthetic symbiosis has been hampered by the lack of model systems. Initially described in extreme deep-sea environments, such as hydrothermal vents, cold seeps, or anoxic basins, these models are logistically and financially difficult to access and are adapted to very high-pressure conditions that are not easily reproducible in the laboratory (Felbeck et al., 1981; Fisher et al., 2007; Dubilier et al., 2008). Consequently, no practical, easy to use model systems have emerged from these environments.

Over the past few decades, interestingly, instances of chemosynthetic symbioses have also been discovered in coastal marine shallow waters. This ease of access has allowed various symbioses to be investigated as potential model systems that could be established under laboratory conditions. Among these, the peritrich ciliate, *Zoothamnium niveum*, found in shallow water habitats, has emerged as a promising candidate due to its global distribution and tractability (Bright et al, 2014), making it less challenging to access in comparison with deep-sea organisms. Lucinid bivalves, which are found in both shallow and deep waters, host chemosynthetic bacteria like *Z. niveum* and thrive in low-oxygen environments. While they represent the only other chemosynthetic symbiosis system with successful laboratory reproduction, their use as a model system is extremely limited, bordering on the unusable (Yuen et al., 2019). The reproduction and maintenance of lucinids in lab settings require complex setups that not only mimic their natural habitats, but also account for their long reproductive cycles (sexual maturity is reached within two to three years), which are impractical for typical laboratory timelines (Berg and Alatalo, 1985; Gros et al., 1996; Gros et al.,1997).

*Z. niveum* colonies, in contrast, have multiple advantages. Their smaller size and potential for rapid growth make them ideal for quick and efficient experimental studies, unlike larger organisms which may take longer to grow. Additionally, because *Z. niveum* reproduces asexually, all its descendants are genetically identical. This consistency reduces variability in experiments, making it easier to design studies and interpret results (Herron et al., 2013; Corliss, 2002). Our work in establishing the *Z. niveum* model system will help the scientific community to advance our knowledge of the physiology, metabolism, behavior, filtration biophysics, and the mutual benefits between the host and symbiont (Vopel et al., 2001; Vopel et al., 2002; Rinke et al., 2007; Kloiber et al., 2009; Røy et al., 2009; Bright et al., 2014; Espada-Hinojosa et al., 2024).

### An affordable, open-source cultivation system

Model systems are essential for studying symbiosis, allowing experimental manipulations and control, and enabling reproducibility. Such systems provide a controlled environment, free from external variables, leading to more definitive insights and allowing detailed examination of interactions and the study of dynamic processes. Zengler et al. (2019) highlighted the importance of simplicity, accessibility, and standardization in microbiome research through the EcoFAB initiative which encouraged the development of standardized and reproducible platforms that help researchers explore complex ecological and symbiotic relationships by manipulating specific variables effectively. The cultivation system we introduce here follows the same principles.

A key feature of our setup involves 3-D printed flow-through chambers that provide the symbiosis with a continuous supply of sulfide and oxygen (Rinke et al., 2007; Bright et al., 2019). While similar setups exist, they are often expensive, custom-made, and not generally available to the scientific community. Lack of access has prevented the community from improving on existing designs, which in turn has only acted to further limit their usability. Most importantly, daily maintenance requirements and need for proximity to the sampling site significantly limits their use in a laboratory setting. To overcome these limitations, we instead leveraged 3D printing technologies and PDMS casting to develop a new, low-cost, open-source cultivation device that can be readily produced in any laboratory with basic equipment.

Current custom setups require daily replacement of the hydrogen sulfide solution, prepared in ultrapure MilliQ® water, provided to the symbiosis via 50 mL syringes and a syringe pump. Maintenance in our system is significantly lowered by substituting the syringes with a 1 L glass bottle equipped with an argon gas bag. This allowed us to increase the volume of the hydrogen sulfide solution and at the same time to prevent its oxidation (oxygen diffuses through the syringe plastic while glass is O2-impermeable). In our setup, the same hydrogen sulfide solution can now be used to feed the cultivation system for several days, allowing the system to self-maintain over weekends, for instance. Secondly, this new system addresses the challenge of achieving homogeneous supply of mingled seawater and sulfidic solutions by i) precisely adjusting the H₂S solution viscosity with NaCl and ii) by implementing a commercially available helical static mixer right before the cultivation chamber (Thakur et al., 2003). These two improvements offer a streamlined, cost-effective approach that enhances reproducibility and scientific research applicability.

### Successful, multi-generational cultivation of Z. niveum under laboratory conditions

Using our system, we demonstrate successful cultivation of *Z. niveum* with minimal maintenance. The settlement rate in our system reached 41.5%, which is comparable to 38.5%; settlement rates reported in Bright et al., (2019). By maintaining stable physico-chemical conditions, we’ve extended colony survival up to 17 days across multiple generations, supporting over 200 colonies, before we stopped culture. This performance surpasses previous systems, where colony lifespans reached 13 days (Rinke et al., 2007). In our system, colony lifespan was 11.9 ± 3.1 days (n=43), compared to 10.9 ± 1.8 days (n=11) reported by Rinke et al. (2007). In Bright et al. (2019), the culture stopped after 7 days. On day 13, Rinke et al. (2007) reported an average of 51.3 ± 9.3 branches (mean ± SD; n=13). In our setup we observed 67.8 ± 4 branches (mean ± SD; N=29), indicating a more robust growth of branches in our experimental setup as compared to Rinke et al. The observed higher growth performance in generation 1 compared to generations 2 and beyond may reflect the initial selection of optimal colonization sites within the chamber. Despite this slight decline, growth rates remained faster than those reported in other studies. Our culture growth averaged 4.4 (± 1) branches per day, compared to 4.4 (± 0.9) branches per day in Bright et al. 2019 by Day 7. This suggests a comparable growth between the two setups. The filter-feeding and contraction behaviors of the host, which are necessary for symbiont access to nutrients, are observed in laboratory cultures just as they are in the natural environment, effectively replicating these ecological interactions (Ott et al., 1998; Ott et al., 2004). We also noted morphological adaptations in the symbionts, with coccoid forms benefiting from better nutrient access due to improved water flow, compared to rod-shaped forms (Vopel et al., 2001; Vopel et al., 2002; Vopel et al., 2005; Rinke et al., 2007). These data demonstrate that that our system more effectively replicates natural conditions compared to previous setups, highlighting its efficacy in fostering natural growth patterns, symbiotic relationships and supporting enhanced colony sustainability.

Managing microbial abundance is crucial in non-axenic culture systems. Controlling the flow rate is an effective method for reducing microbial abundance, confirming its critical role in maintaining microbial balance (Provoost et al., 2022). Our system introduces seawater containing environmental bacteria, which serve as a food source for *Z. niveum,* in a highly controlled manner. This setup reflects a natural but controlled exposure to microbial communities, important for maintaining a stable culture environment without resorting to strictly axenic conditions. This approach is built on the understanding of the established challenges and proven techniques in microbial management within culture systems developed in Lewis et al., 1988.

### Challenges and prospects for characterizing Zoothamnium niveum chemosynthetic symbiosis

Despite successful cultivation, a few challenges remain. For instance, while we know our current culture is not monoxenic, it remains uncertain whether achieving a monoxenic culture will be possible in the future. The median MA in culture 3 was 4×10^5^, and in culture 4 it was 2.6×10^5^, yielding a combined median of 2.9×10^5^. This is approximately 2.9 times lower than that reported by Bright et al. (2019) which held a MA of 7.9×10^5^ for experiment 68 and 7.5×10^5^ for experiment 75. This indicates that our cultures contained two to three times fewer microbes. This is likely due to slower flow rates and finer seawater filtration (1.2 μm) in our system, which we use to remove more contaminants and prevent biofilm buildup. This also enhances stability and replicability compared to earlier setups like those by Bright et al. with a 32 µm filtered seawater.

Our proposed set-up is well equipped to study various aspects of chemosynthetic symbioses. Robust growth of *Z. niveum* and its longevity allow sample collection at timed intervals to study various growth phases. The apparatus is equipped with a glass slide on top allowing for the placement of a dissecting microscope, facilitating direct observation and imaging. This allows developmental changes and the colony’s dynamics to be tracked in real time, as well as for conducting time-lapse photography to image the cultivated specimen’s behaviors such as its contractions, growth and perhaps new features never seen before. For instance, it allowed us to observe the release of a coat, which is a new circular/ovoid structure expelled out of the terminal zooid with a rough texture suggesting bacterial involvement, confirmed through confocal observation. These structures have not been reported in previous studies on *Z. niveum* or other ciliates and their exact roles are still under investigation.

We envision the use of the *Z. niveum* system by the scientific community for detailed molecular insights into the mechanisms of symbiosis, as well as detailed genomic and proteomic characterization of both host and symbiont. Transcriptomic studies on *Z. niveum* will reveal changes in gene expression in response to environmental variations and help identify key genes involved in symbiosis. Genomic studies of *Z. niveum* and its symbiont *Thiobius zoothamnicola* will uncover potential genomic adaptations that could explain the ecological success of this symbiont, on a sessile host across varied environments all over the globe. Other techniques such Cryo-electron microscopy, made possible given the ready availability of samples, can reveal how physical structures support their chemosynthetic lifestyles, offering insights into cellular mechanisms that underpin symbiosis at a resolution not achievable with other microscopy techniques (Tanaka et al., 2019).

## Acknowledgments

We are thankful to the electron microscopy facility C3MAG “Centre Commun de Caractérisation des Matériaux des Antilles et de la Guyane” in Guadeloupe, F.W.I., which is supported by The European Regional Development Fund, the Regional Council of Guadeloupe, and the French Research Department. We are thankful to Janey Lee for technical support on this project.

The work by J.-M.V. (proposal: 10.46936/10.25585/60001074) conducted by the US Department of Energy Joint Genome Institute (https://ror.org/04xm1d337), a Department of Energy (DOE) Office of Science User Facility, is supported by the Office of Science of the DOE operated under contract no. DE-AC02-05CH11231. P.-E.C. is supported by a grant from the Collectivité Territoriale de Martinique (convention number 31.03.2022-0236). This work was supported by the Gordon and Betty Moore Foundation (grant GBMF9340 to J.-M.V., S.V.D., and P.-E.C.). We would also like to thank Erik Beahrs for his assistance in designing the 3D model of the mold and to the Makerspace at the UCSB library for their support.

