## supplementary material for "A novel open-source cultivation system helps establish the first full cycle chemosynthetic symbiosis model system involving the giant ciliate *Zoothamnium niveum*"

[Supplementary movie 1](https://drive.google.com/drive/u/0/folders/1Y1AUum__SEJ5egnggr3C2iwVoawgkCRo) - Timelapse photography capturing the first 10 hours of growth of a colony of *Z. niveum.* The timelapse starts right after the swarmer settled on a side of the chamber, differentiated into a terminal zooid and started to produce a stalk. During the first 1h40min, the stalk grows without symbionts, with the terminal zooids undergoing rotations. After that time, there is a pause in the stalk elongation. The terminal zooid changes its behaviour and after approximately 50 min it sheds a coat., the growth resumes, with the symbionts now covering the entire colony, including the stalk.

[Supplementary movie 2](https://drive.google.com/drive/u/0/folders/18Y6XOUqucfN99xnAW3pS1omzcjGSlOn0) - Timelapse photography, 105 minutes long, of 5 *Zoothamnium niveum* colonies after 8 days of cultivation showing their contraction behaviour. Note that the cultivation setup is well suited for live imaging of the symbiosis and using a simple dissecting microscope and with a camera, quantitative data can easily be produced. Here for instance colonies from left to right contract at a frequency of 0.29, 0.26, 0.3, 0.28, 0.29 contractions/min respectively with a mean 0.284±0.0152.

Supplementary table 1 - Documented Sampling Sites and Environmental Contexts of *Zoothamnium niveum*. The table lists all known sampling locations for *Zoothamnium niveum*, detailing the geographic sites, coordinates, depths, and environmental substrates where specimens have been collected.

| **Number** | **Localisation** | **GPS point** | **Depth (meters)** | **Environment** | **Substrate** | **Reference** |
| --- | --- | --- | --- | --- | --- | --- |
| 1 | Twin Cays Island |  |  | Shallow water, tidal channel cut into mangrove peat | rocks | [Rinke et al., 2006](https://www.frontiersin.org/articles/10.3389/fmicb.2014.00145/full#B66) |
| 2 | Florida keys, Gulf of Mexico |  |  |  | rotting plant material | [Bauer-Nebelsick et al., 1996a](https://www.frontiersin.org/articles/10.3389/fmicb.2014.00145/full#B2) |
| 3 | Guadeloupe, French West Indies | 16°N, 61.5°W | 0.5-1.5 m | Shallow water, on mangrove sediment | sunken wood and leaf debris | [Laurent et al., 2009](https://www.frontiersin.org/articles/10.3389/fmicb.2014.00145/full#B43), [2013](https://www.frontiersin.org/articles/10.3389/fmicb.2014.00145/full#B44) |
| 4 | Canary Islands, Lanzarote | 28°55'05''N -13°39'36''W | 6 m | On the sides of a rocky depression | rocks | [Wirtz and Debelius, 2003](https://www.frontiersin.org/articles/10.3389/fmicb.2014.00145/full#B85) |
| 5 | Corsica, France |  |  | Near *Sargassum* spp. debris accumulation | rocks | Bright et al., 2014 |
| 6 | Adriatic sea |  |  |  | sunken woods | Bright et al., 2014 |
| 7 | Red sea |  |  |  | rotting plant material | Bright et al., 2014 |
| 8 | Tokyo bay |  |  | Deployed whale fall | bones | [Kawato et al., 2010](https://www.frontiersin.org/articles/10.3389/fmicb.2014.00145/full#B41) |
| 9 | White Point Beach California |  |  | Tidal pool close to coastal hydrothermal vents |  | Volland J-M., personal observation |
| 10 | Florida keys, Indian River Lagoon; East coast | 27°25'11.78"N, 80°16'30.575"W |  |  | wooden debris | Indian River Lagoon Species Inventory; Clamp and Williams, 2006 |
| 11 | Madeira Island |  |  | At the bottom of two large pools, Praia Piscina |  | Wirtz unpublished |
| 12 | São Tomé Island | 0°01´25´´ N, 6°30´45´´ E | 1.5 m |  | rotting coconuts | Wirtz, 2018 |
| 13 | Gabon, coastal area | 0°36´64´´N, 9°18´43´´E | 1 m |  |  | Wirtz & Serval-Roquefort., 2020 |
| 14 | Giglio Island, Western Mediterranean | 42°22'01''N - 10°53'33''E | 30 m | On sandy bottom | rocky tree trunk | Wirtz, 2008 |
| 15 | Cyprus Island | 34°57'45''N,34°04'26''E |  | In a cave, close to cracks in the wall | rock wall | Wirtz, 2008 |

Supplementary Table 2 - comparison of the genome statistics of *Zoothamnium niveum*’s symbiont*, Ca.* Thiobius zoothamnicola, isolated from California and from the Caribeean (Belize and Guadeloupe). The three draft genomes have similar statistics. GC Content

Contiguity and Quality of Assemblies: The number of contigs and the N50 values provide metrics for contiguity and quality of the genomic assemblies. All locations demonstrate high-quality assemblies with similar numbers of contigs (46 for Belize and Guadeloupe, and 45 for California) and substantial N50 values, indicating that the assemblies are well suitable for genomic investigations.

Completeness and Contamination: Completeness of the genomes is high across all samples, with Belize and Guadeloupe showing equal completeness estimates at 96.0%, while California shows slightly lower estimates at 85.5%. Heterogeneity: All samples exhibit zero percent estimated heterogeneity, suggesting that the assemblies represent single strains without significant intra-sample genetic diversity.

|  | *Ca*. Thiobius zoothamnicola | | |
| --- | --- | --- | --- |
|  | Belize | Guadeloupe | California |
| Assembly size (bp) | 2,381,364 | 2,369,374 | 2,151,318 |
| GC content (%) | 49,40 | 49,60 | 49,52 |
| nbr. of contigs | 46 | 46 | 45 |
| N50 (bp) | 98,217 | 73,149 | 69,285 |
| Completeness (%) | 96.0 | 96.0 | 85.5 |
| Contamination (%) | 0.1 | 0.4 | 0.0 |
| Heterogeneity (%) | 0.0 | 0.0 | 0.0 |

Supplementary table 3 - Information about the seven genome bins obtained from the *Zoothamnium niveum* metagenome, including the Genome Taxonomy Database taxonomic assignment, the total size of the bin assembly, the number of contigs, the GC percentage, the estimated completeness, the estimated contamination level, the number of predicted genes, and the average protein length. The first bin is a taxonomic assignment to the genus *Aliishimia*. Bin 6, and 7 do not have a taxonomic assignment with a completeness below 0.1%. Bin number 3 contains eukaryotic sequences and corresponds to the ciliate host. Bins 3 and 4 are gammaproteobacterial draft genomes. Only one of these bins unambiguously emerged as the host's symbiont with an ANI of 96.7% compared to the published genome *Ca*. Thiobios zoothamnicola.

| Bin# | Taxonomic Assignment | species | size (bp) | contig# | GC (%) | est. Comp. % | cont. % | predicted genes | Aver. Prot length (aa) |
| --- | --- | --- | --- | --- | --- | --- | --- | --- | --- |
| 1 | f__Rhodobacteraceae;g__Aliishimia |  | 2,168,570 | 230 | 54.5 | 79.7 | 0.1 | 2,241 | 279 |
| 2 | none |  | 426,229 | 81 | 36.5 | 0.9 | 0.0 | 531 | 83 |
| 3 | Eukaryote (Blast n with selected genes) | *Z. niveum* (host) | 38,645,428 | 697 | 33.9 | 54.8 | 62.8 | 56,610 | 85 |
| 4 | f__Thiotrichaceae |  | 2,946,997 | 69 | 40.8 | 94.3 | 1.1 | 2,534 | 337 |
| 5 | f__Sedimenticolaceae;g__VMDJ01 | *T. zoothamnicola* (symbiont) | 2,151,318 | 45 | 49.5 | 85.5 | 0.0 | 2,137 | 303 |
| 6 | none |  | 204,369 | 50 | 33.3 | 0.0 | 0.0 | 204 | 75 |
| 7 | none |  | 546,988 | 87 | 34.2 | 0.0 | 0.0 | 307 | 86 |

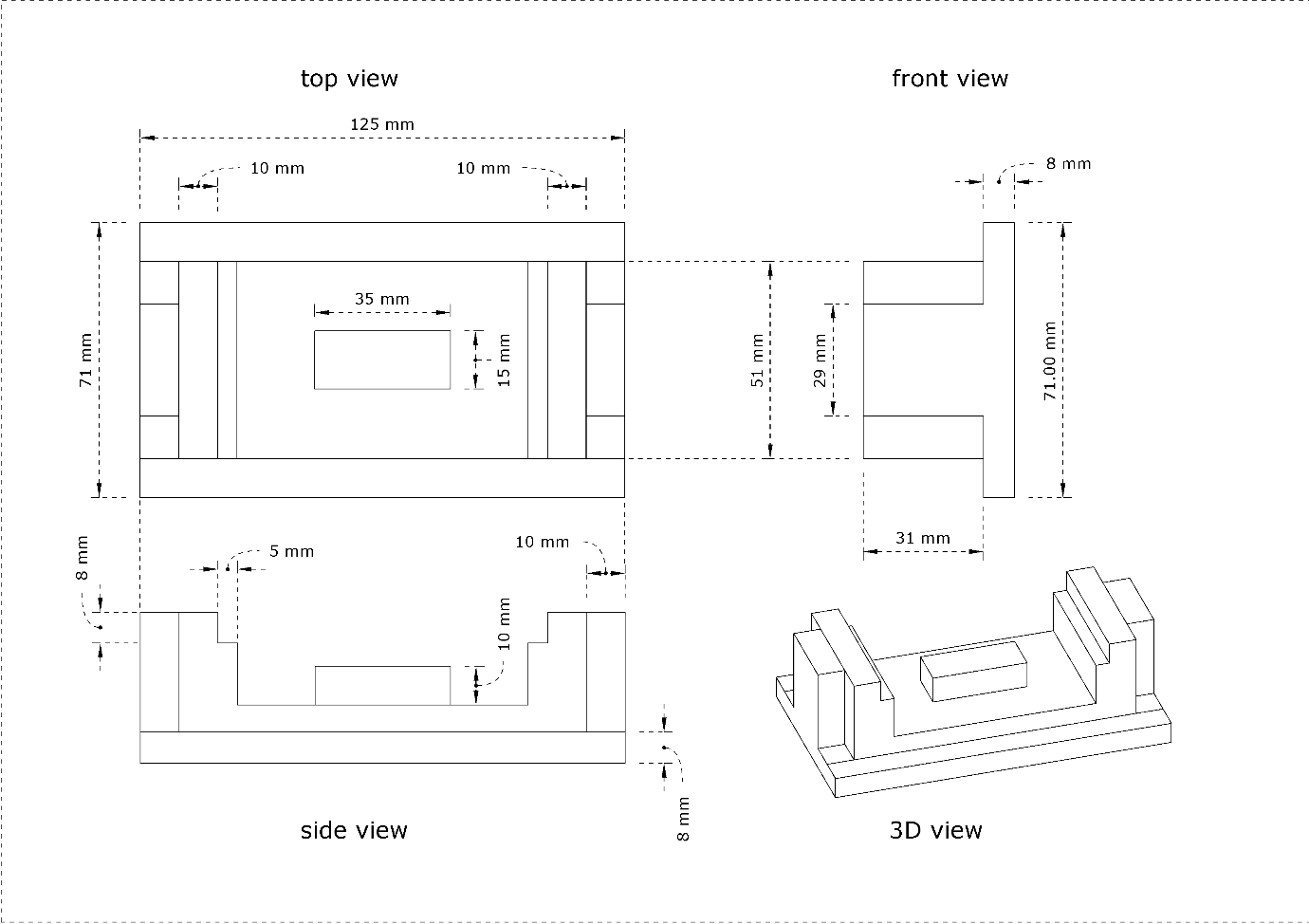

Figure S1 – Technical drawings of the central part of the mold. Top view, front view, side view, and 3D view of the central section of the mold used for casting the PDMS cultivation chamber. The stl file of this 3D model is available for download at https://github.com/jvolland/Zoothamnium-Cultivation-System.

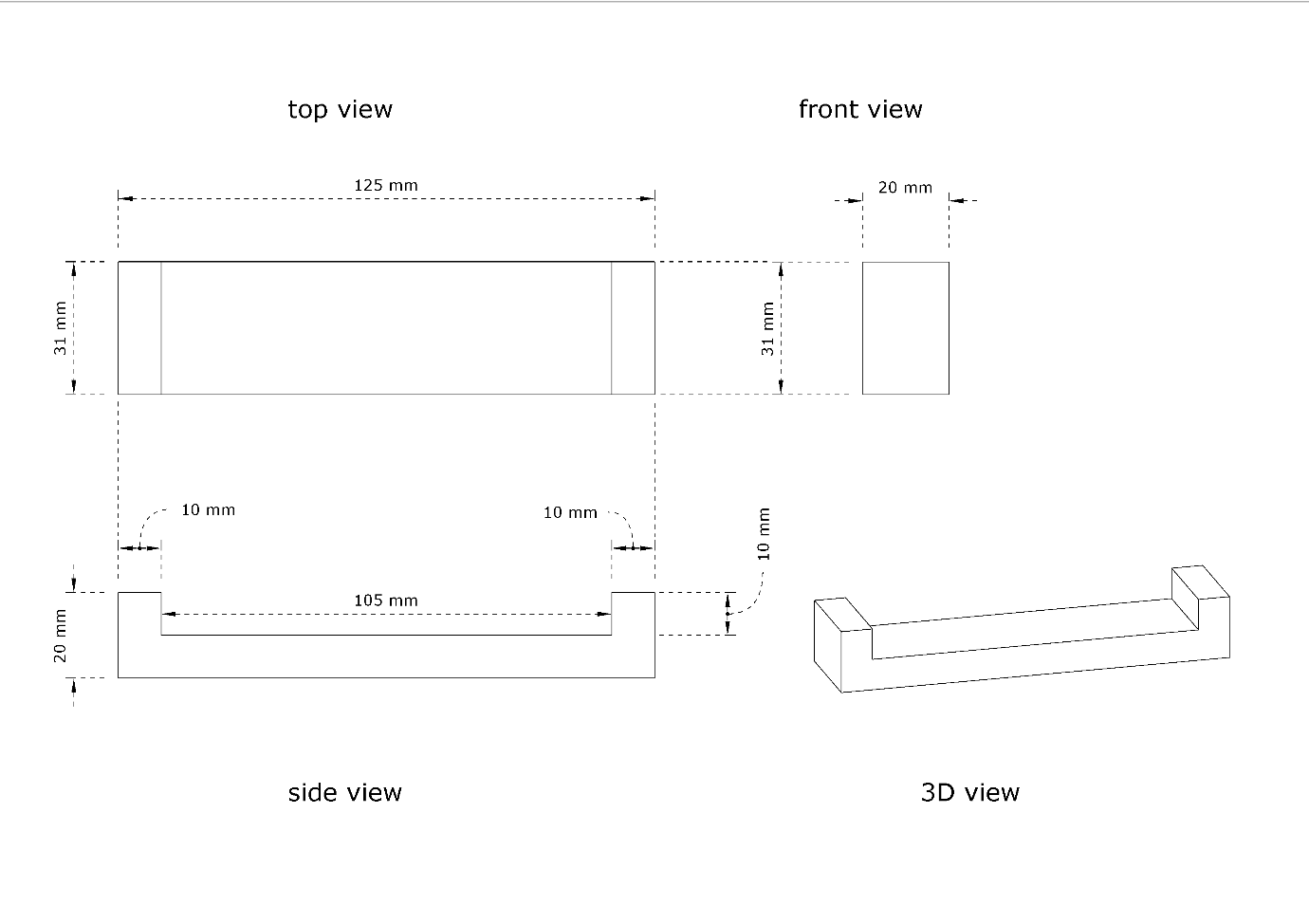

Figure S2 – Technical drawings of the side part of the mold. Top view, front view, side view, and 3D view of the side section of the mold used for casting the PDMS cultivation chamber. This side part is designed to close the two long edges of the central piece of the mold presented in Figure S1. The stl file of this 3D model is available for download at https://github.com/jvolland/Zoothamnium-Cultivation-System.

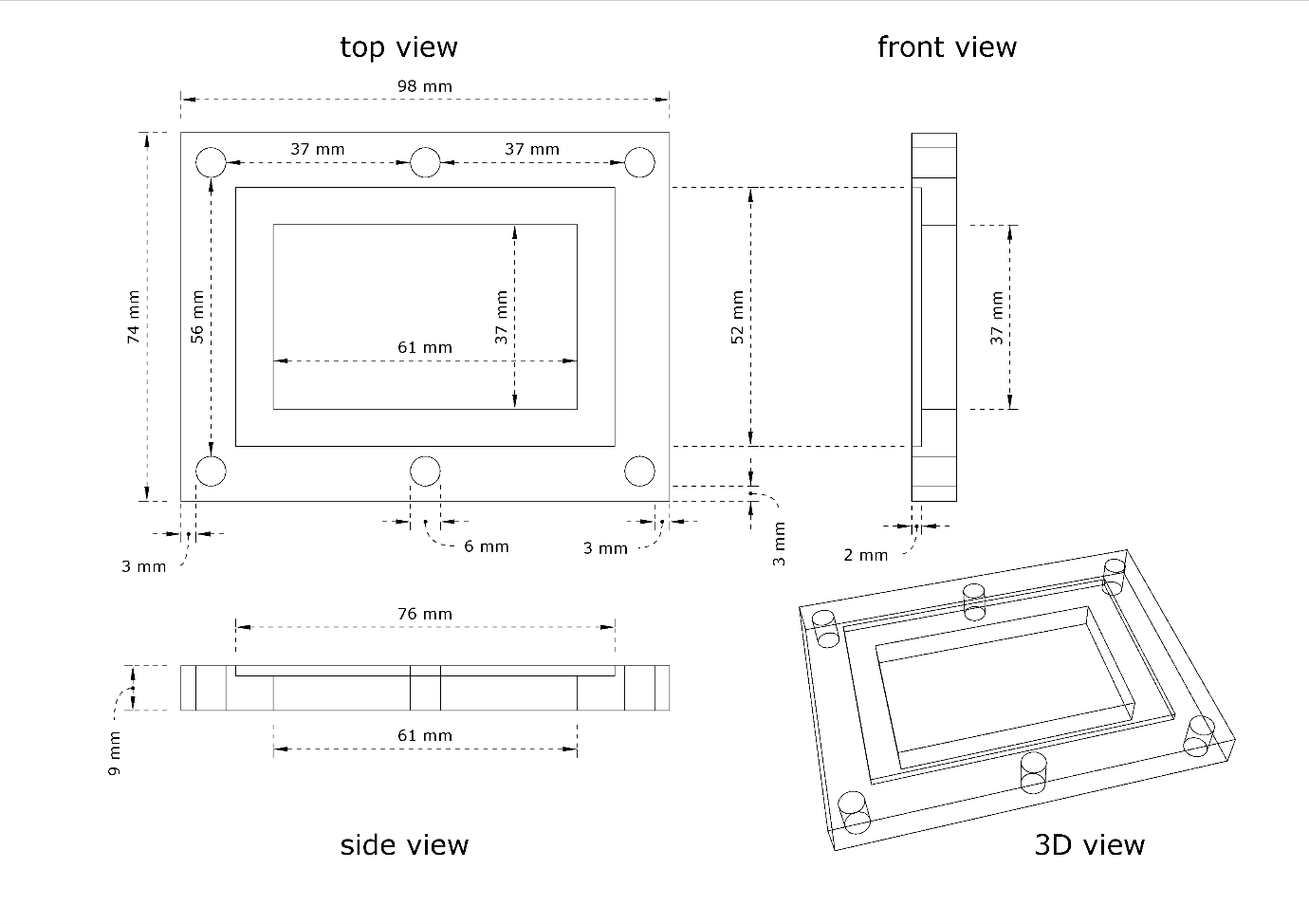
Figure S3 – Technical drawing of the cultivation chamber frame. Top view, front view, side view, and 3D view of the frame used for assembling the cultivation chamber. The frame secures the glass slide onto the PDMS body and maintains the alignment of the components. The top view shows the placement of six bolt holes that are used to hold the structure together, ensuring a tight and leak-free seal. The stl file of this 3D model is available for download at <https://github.com/jvolland/Zoothamnium-Cultivation-System>.

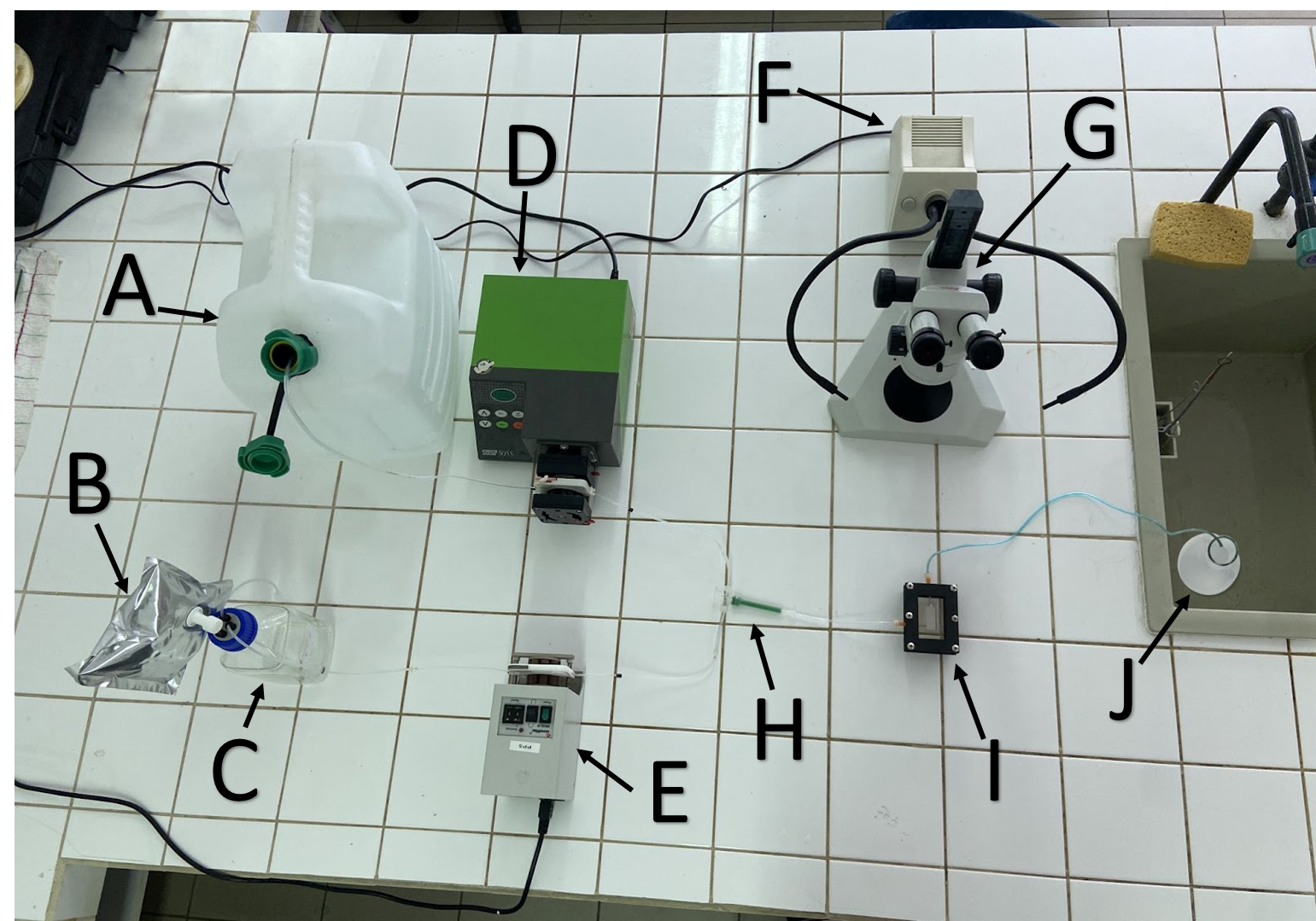

Figure S4 - Overview of the culture system. (A) filtered sea water tank; (B) gas bag filled with argon; (C) anoxic sulfide solution with 20gr/L NaCl; (D) peristaltic pump for sea water; (E) peristaltic pump for sulfidic water; (F) light source; (G) dissection microscope; (H) static mixer; (I) culture chamber; (J) outlet water collection.

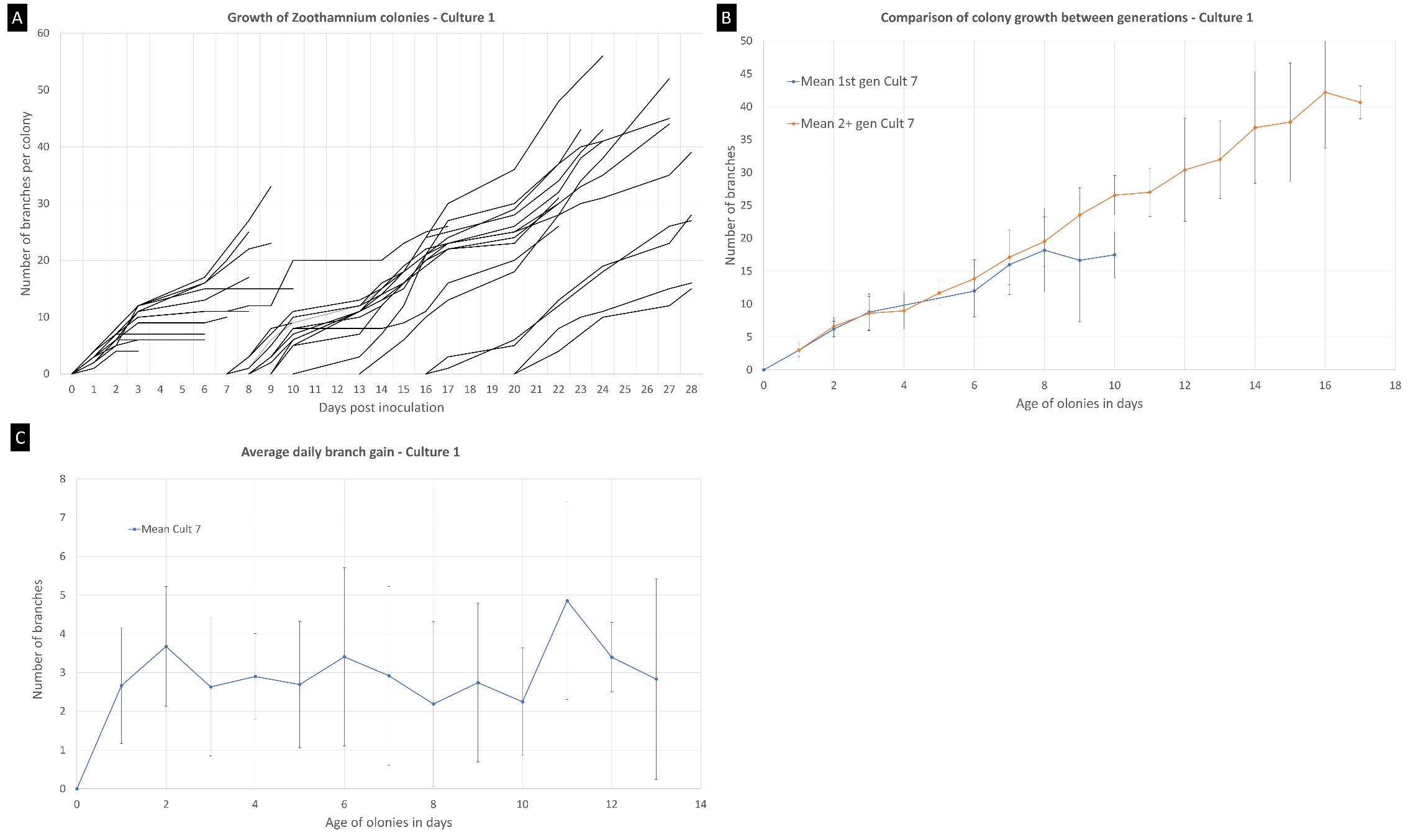

Figure S5 - Growth of *Zoothamnium niveum* culture #1. (A) Growth of randomly selected individual colonies. The second generation of colonies start to grow after 7 days.; (B) Comparison of colony growth between the first generation (n=13) and the following generations (n=16); (C) Average branch gain per day measured on our culture (n=29).

B

C

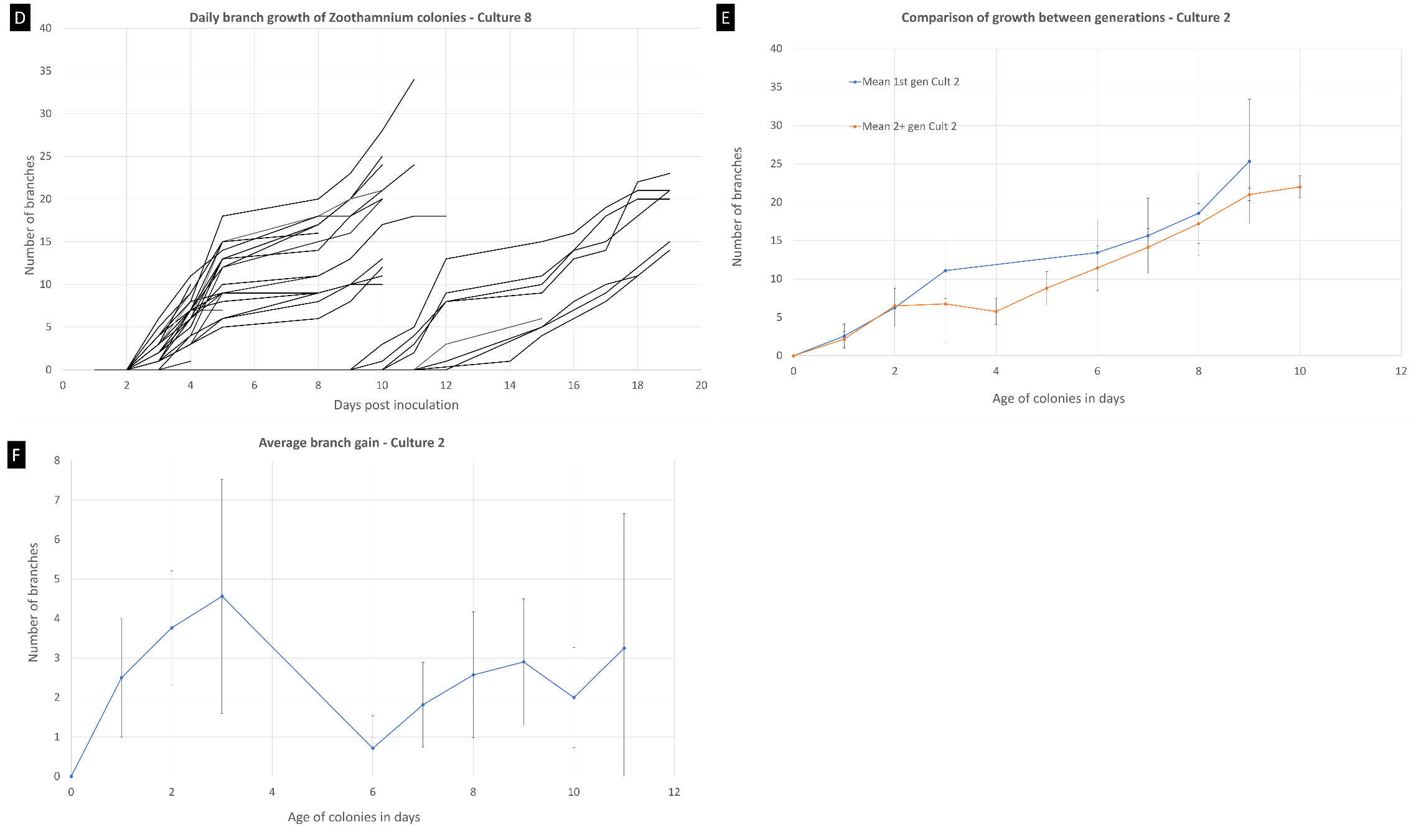

Figure S6 - Growth of *Zoothamnium niveum* culture #2. (A) Growth of randomly selected individual colonies. The second generation of colonies start to grow after 9 days.; (B) Comparison of colony growth between the first generation (n=18) and the following generations (n=10); (C) Average branch gain per day measured on our culture (n=28).

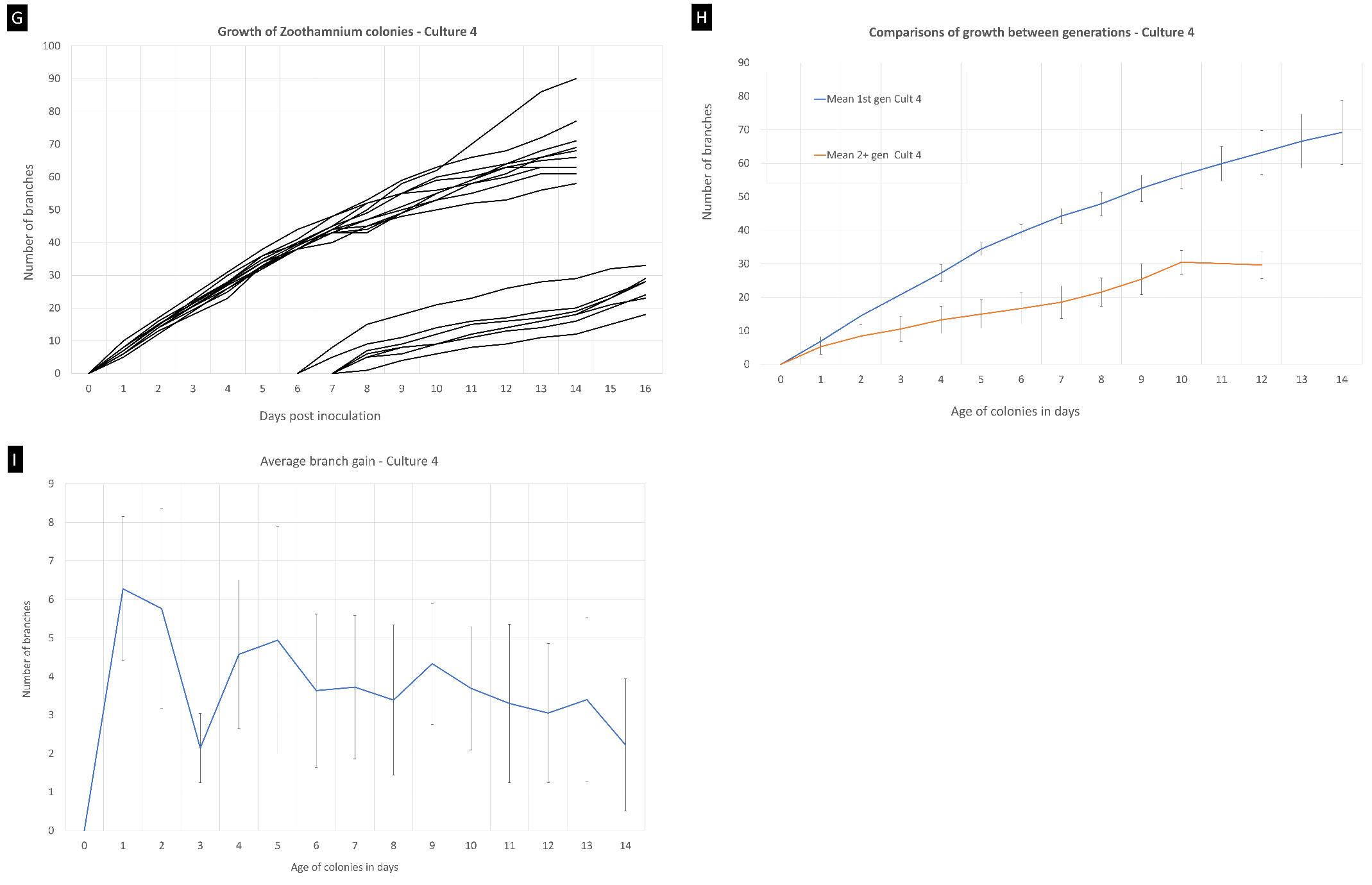

Figure S7 - Growth of *Zoothamnium niveum* culture #1. (A) Growth of randomly selected individual colonies. The second generation of colonies start to grow after 7 days.; (B) Comparison of colony growth between the first generation (n=13) and the following generations (n=16); (C) Average branch gain per day measured on our culture (n=29).

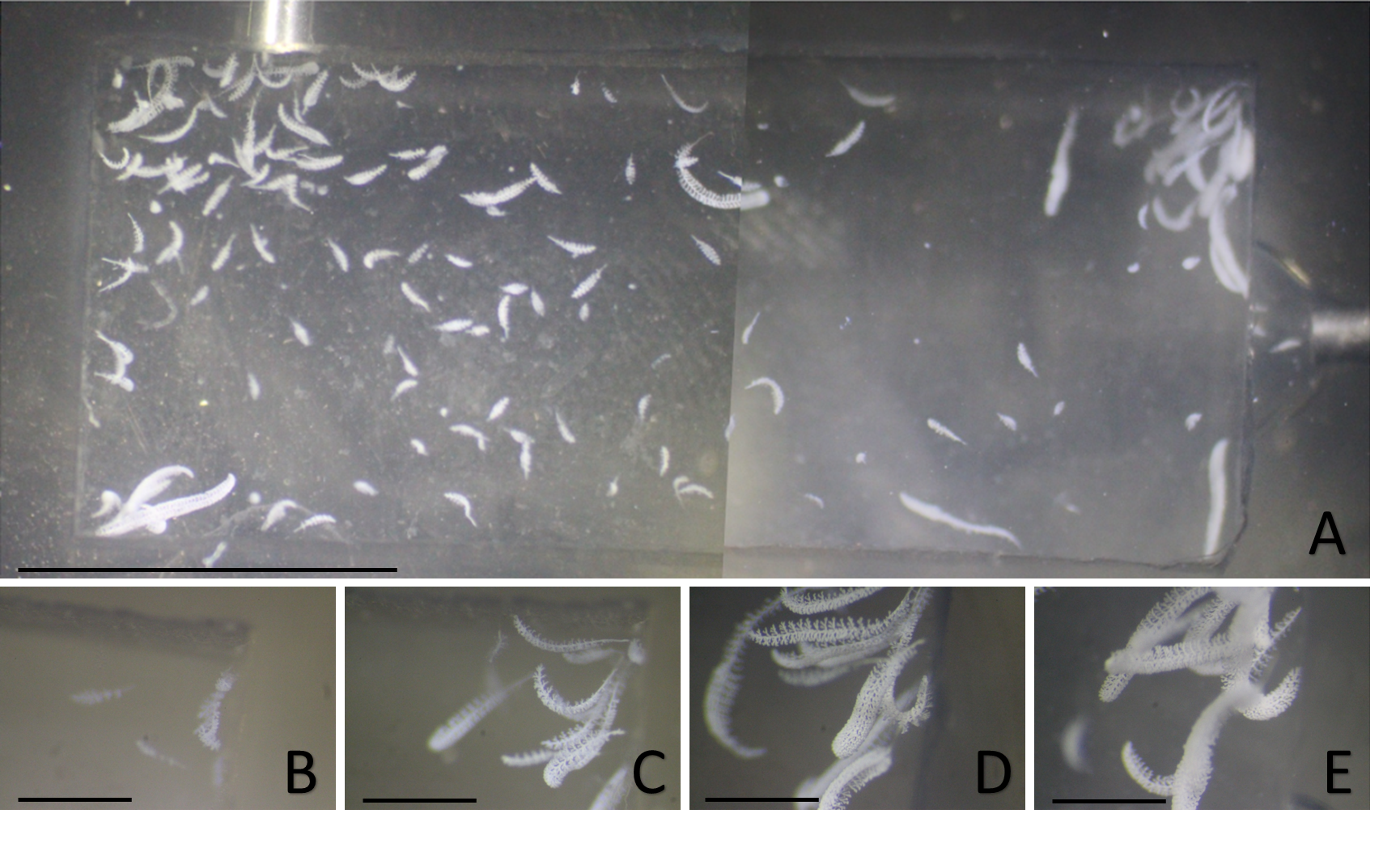
Figure S8 – Overview of culture #3. (A) Photo-montage of the whole culture chamber 15 days after incubation; scale bar = 1 cm. The detailed view of the top right corner is given at higher magnification and shows the growth of *Zoothamnium niveum* colonies after 1 day (B), after 5 days (C), after 10 days (D), and after 15 days of laboratory culture; scale bars = 0.25 cm.

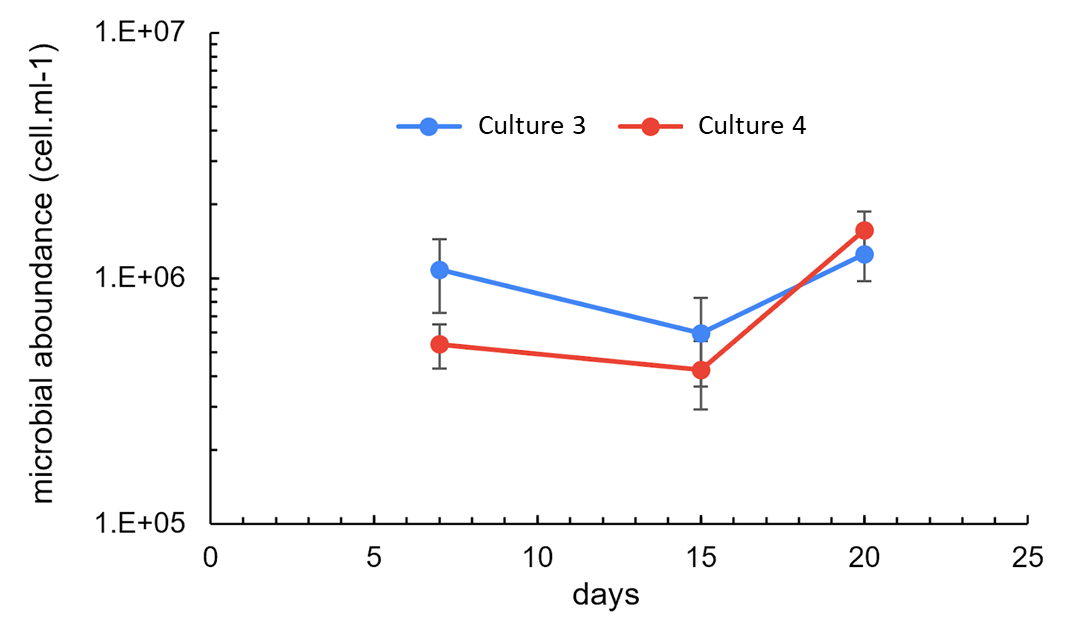

Figure S9 – Monitoring of microbial abundance in the outlet water of cultures 3 and 4.

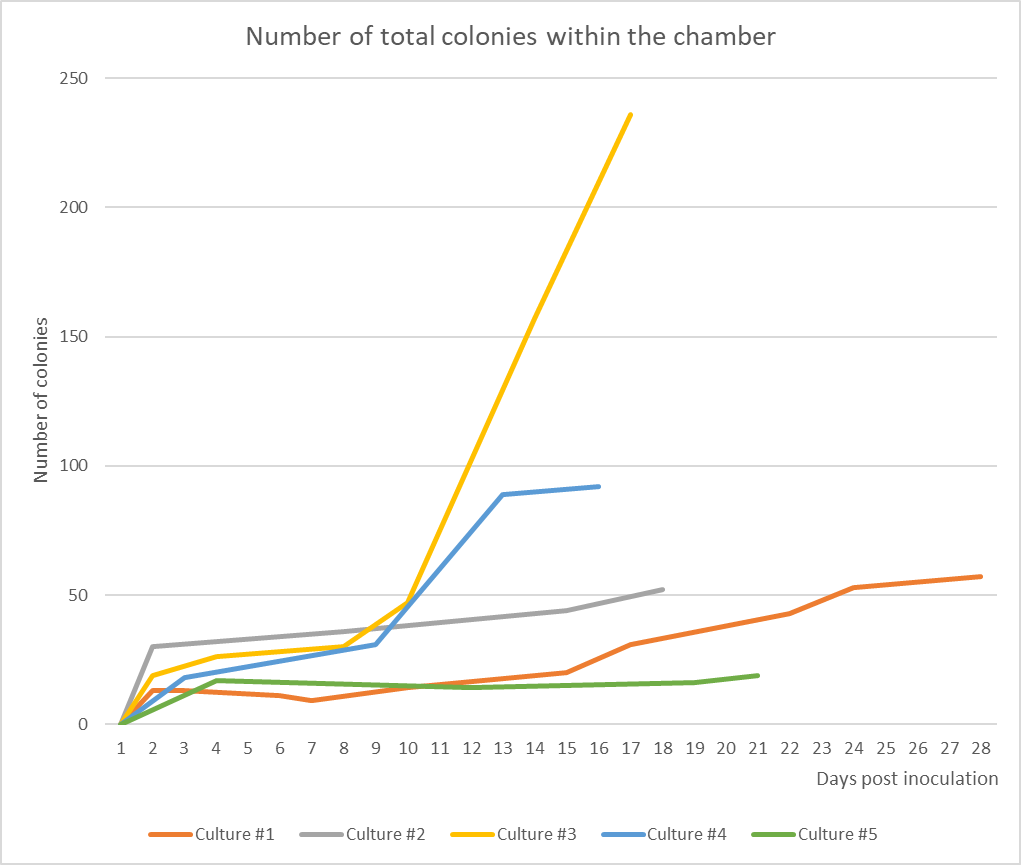

Figure S10 – Evolution of the number of colonies within cultures. The number of swarmers injected in the chambers as inoculum and the percentage of settlements were as followed: culture #1: 38 swarmers with 34% settlement; culture #2: 68 swarmers with 44% settlement; culture #3: 82 swarmers 41% settlement; culture #4: 61 swarmers 51% settlement; culture 5: 19 swarmers.

Supplementary data 1 -*Ca*. Thiobius zoothamnicola’s 16S rRNA gene sequence (sampled from White Point Beach, CA, USA).

>WPB_14_Thiobius_zoothamnicola16S

GAACTGAAGAGTTTGATCCTGGCTCAGATTGAACGCTGGCGGCATGCTTAACACATGCAAGTCGAACGCGAAAGCGCTTTCGGGCGTTAGTAGAGTGGCGGACGGGTGAGTAACGCGTGGGAATCTACCTGATAGTGGGGGATAACTCGGGGAAACTCGAGCTAATACCGCATACGCCCTACGGGGGAAAGACTTCGGTCACTATCAGATGAGCTCGCGTCAGATTAGCTTGTTGGTGAGGTAATGGCTCACCAAGGCAACGATCTGTAGCTGGTCTGAGAGGACGAACAGCCACACTGGGACTGAGACACGGCCCAGACTCCTACGGGAGGCAGCAGTGAGGAATATTGCACAATGGGGGAAACCCTGATGCAGCAATGCCGCGTGTGTGAAGAAGGCCTGCGGGTTGTAAAGCACTTTCAGTTGGGAAGATAATGACGTTACCAACAGAAGAAGCACCGGCTAACTCCGTGCCAGCAGCCGCGGTAATACGGAGGGTGCAAGCGTTAATCGGAATTACTGGGCGTAAAGCGCGCGTAGGCGGCTAGGTCAGTCAGATGTGAAATCCCCAGGCTCAACCTGGGAATTGCATTTGATACTGCTTGGCTAGAGTATGGTAGAGGGTGGTGGAATTCCAGGTGTAGCGGTGAAATGCGTAGATATCTGGAGGAACATCAGTGGCGAAGGCGGCCACCTGGATCAATACTGACGCTGAAGTGCGAAAGCGTGGGGAGCAAACAGGATTAGATACCCTGGTAGTCCACGCCGTAAACGATGTCAACTAGCCGTTGGGTTTTTATAAATTTAGTGGCGTAGCTAACGCGATAAGTTGACCGCCTGGGGAGTACGGCCGCAAGGTTAAAACTCAAAGGAATTGACGGGGGCCCGCACAAGCGGTGGAGCATGTGGTTTAATTCGATGCAACGCGAAGAACCTTACCAGTTCTTGACATCCAGTGAACTTTCCAGAGATGGATTGGTGCCTTCGGGAACACTGTGACAGGTGCTGCATGGCTGTCGTCAGCTCGTGTCGTGAGATGTTGGGTTAAGTCCCGTAACGAGCGCAACCCTTGTCCTTAGTTGCCAGCACTTCGGGTGGGAACTCTAAGGAGACTGCCGGTGACAAACCGGAGGAAGGTGGGGATGACGTCAAGTCATCATGGCCCTTATGAACTGGGCTACACACGTGCTACAATGGACGGTACAGAGGGCTGCAAACCCGCGAGGGGGAGCTAATCCCAAAAAACCGTTCGTAGTCCGGATTGCAGTCTGCAACTCGACTGCATGAAGTCGGAATCGCTAGTAATCGCAGATCAGAATGCTGCGGTGAATACGTTCCCGGGCCTTGTACACACCGCCCGTCACACCATGGGAGTGGGTTGCAAAAGAAGTGGGTAGCTTAATAATGGGCGCTCACCACTTTGTGATTCATGACTGGGGTGAAGTCGTAACAAGGTAGCCGTAGGGGAACCTGCGGCTGGATCACCTCCTTT

Supplementary data 2 - *Zoothamnium niveum*’s 18S rRNA gene sequence (sampled from White Point Beach, CA, USA).

>Zoothamnium_niveum_WPB_partial18S

AATCTGGTTGATCCTGCCAGTAGTCATATGCTTGTCTCAAAGATTAAGCCATGCATGTGTAAGTATAAGTATTATACGGCGAAACTGCGAATGGCTCATTAAATCAGTTATAATTTATTTGATAATCGAAAGTTACATGGATAACCGTGACAAATTACAGCTAATACATGCAGTCAGACCCGGTCCAAAGGTCGTAATTATTAGTATTAAACCAATTCCTTCGGGAGTGTGATGAATCATAGTAATCGAACGAATCGCTAATCTTGCGATAAATCATTCAAGTTTCTGCCCTATCAGCTTTGGATGGTAGTGTATTGGACTACCATGGCAGTCACGGGTAACGGAGAATTAGGGTTCGATTCCGGAGAGGGAGCCTGAGAAACGGCTACCACATCTACGGAAGGCAGCAGGCGCGAAAATTGCCCAATCCTGACACAGGGAGGCAGTGACAAGAAATAACAACTCTCGGTTTCCGAGAAGTGTAATGAGGATAATTTACAAACCTTACCGAAAGCAATTGGAGGGCAAGTCTGGTGCCAGCAGCCGCGGTAATTCCAGCTCCAATAGCGTATATTAAAGTTGTTGCAGTTAAAAAGCTCGTAGTTGAAATTCTGATTGTATTGGGCCTCAGCACTCGATGCGTAGGTACCAGCAGTCATCCGCTTGCAAACATATGTTCTTCCTTAAACAGAAGACTATGTGAGTAAGCATTTTACCTTGAGAAAAACAGAGTGTTCCAGGCAGGTTTGTCCGGAATGCATTAGCATGGAATAATAGAATATGACTGGAGTCTATTTATTGGTTTGAGACTACAGTAATGATTAATAGGAACAATCGGGGGCATTGGTACTTGTCAGTCAGAGGTGAAATTCTAGGATTTGACAAAGACTAACAAATGCGAAAGCATTTGCCAAGGATGTTTTCATTAATCAAGAACGAAAGTTAGGGGATCAAAGACGATCAGATACCGTCCTAGTCTTAACTATAAACCATACCGACTCGGATTCAGATGAATAATAAAGTTCATTTGGGACCGTAGGAGAAATCAAAGTCTTTGGGTTCTGGGGGGAGTATGGTCGCAAGGCTGAAACTTAAAGGAATTGACGGTTTTGCACCACCATGGAGTGGAGCCTGCGGCTTAATTTGACTCAACACTGGGAAACTCATCAGGGCAAGAAGATTTTAGGATTGACAGATTGAGAGTTCTTTCTTGATTGGTCTAGTGGTGGTGCATGGCCGTTCTTAGTTGGTGGAGTGATTTGTCTGGTTAATTCCGTTAACGAACGAGACCTTAACCTGCTAACTAGTATTGCTATCCACAATGGCAATTACTTCTTAGAGGGACTATGTGATGCAATCACATGGAAGTTTGAGGCAATAACAGGTCTGTGATGCCCTTAGATGTCCTGAGCTGCACGCGCGCTACAATGACACATTCAACGAGCATTTCCTGATCCGAAAGGATTTGGGTAATCTTTTCAGTATGTGTCGTGCTAGGGATAGATCTTTGTAATTGTGGATCTTGAACGAGGAATTCCTAGTAAGCACAAGTCATCAGCTTGTGCTGATTACGTCCCTGCAAAATGTACACACCGCCCGTCGCTGTTACCGATTGAGTGCTCAGGTGAACCTTCTTGATAGTGTCAAAACTAAAAATTAAGTAAACCTTAGCACTTAGAGGAAACAAAAGTCGTAACAAGGTTTCCGTAGGTGAACCTGCGGAAGGATCATTT
